# Divided Attention in Perception: A Unified Analysis of Dual-Task Deficits and Congruency Effects

**DOI:** 10.1101/2020.01.23.917492

**Authors:** John Palmer, Alex L. White, Cathleen M. Moore, Geoffrey M. Boynton

## Abstract

How well can one perceive simultaneous stimuli at two widely spaced visual locations? Are the stimuli processed independently? If not, does the dependency affect perception, disrupt signals in later stages, or both? To address these questions, we measured effects of divided attention using a dual-task paradigm with stimuli presented in noise on either side of fixation. This paradigm was applied to detecting Gabor patches and to the semantic categorization of words. We measured *dual-task deficits* which are a decline in mean performance for a dual task compared to a single task. There was such a deficit for categorizing two words but relatively little deficit for detecting two Gabors. We also measured *congruency effects* which are when performance at one location depends on whether the stimulus at the other location requires the same response. There was such a congruency effect for detecting two Gabors but relatively little congruency effect for categorizing two words. Further experiments were consistent with the dual-task deficit in word categorization being perceptual, but the congruency effect in Gabor detection being due to later processes. Results of additional experiments showed that the congruency effect was consistent with either graded selection errors or with all-or-none selection followed by graded interactive processing. To answer our opening question: for Gabor detection, perceptual processes were largely independent but later processes caused congruency effects; for word categorization, perceptual processes had capacity limits but even in combination with later processes caused relatively little congruency effects. In summary, there was evidence for two different kinds of dependency. Such complementary dependencies are inconsistent with theories of divided attention that depend on a single dependency such as a single resource or single source of interactive processing.

---

To study divided attention in perception, one can ask if multiple stimuli can be perceived as well as a single stimulus. Some theories pose a simple answer to this question based on a single dependency between the processing of each stimulus. For example, a single bottleneck theory predicts divided attention effects for a combination of stimuli and judgment that requires processing of each stimulus after the bottleneck (Broadbent, 1958). In contrast, other theories allow for multiple dependencies between the processing of each stimulus. For example, there might be both capacity limits for perceiving objects (Duncan, 1984) and interactive processing of their representations in memory, decision and response (response competition, Eriksen & Schultz, 1979). To address these alternative theories, we analyze two measures of divided attention effects for a pair of tasks chosen to reveal the diversity of observed effects.

## Domain of Study

Our overarching goal is to understand divided attention in perception. To that end, we focus a particular kind of dual-task paradigms and measure both dual-task deficits and congruency effects. In the following few paragraphs, we introduce this kind of dual-task paradigm, the two measurements, and the issue of isolating perceptual processes.

### Accuracy dual tasks with one kind of judgment

In the dual-task paradigm, one varies the number of relevant stimuli and requires a separate judgment for each stimulus. A decline in performance with two judgments (dual-task deficit) is a divided attention effect. Such tasks are sometimes called concurrent tasks and an introductory review can be found in Sperling and Dosher (1986). We focus on dual tasks that involve separate but similar judgments of two stimuli. For example, one might have to detect a light to the left of fixation, and detect a light to the right of fixation. Or, one might have to identify a word to the left of fixation, and identify a word to the right of fixation. Typically the stimuli are simultaneously and briefly presented to challenge perception. The task is not speeded to prevent dependencies due to preparing and executing speeded responses. Thus, this paradigm is distinct from the large world of speeded dual tasks which are also referred to as the psychological refractory period (PRP) paradigm. There is a bountiful literature on divided attention effects in the speeded paradigm that concentrate on the consequences of divided attention for decision and response processes (e.g. Welford, 1952; Pashler & Johnston, 1998). Using accuracy dual tasks helps shift the focus to perceptual processes. We also restrict ourselves to testing the same perceptual judgment for both tasks. This restriction focuses the work on understanding divided attention for a particular judgment rather than for combinations of judgments.

### Two measurements

In such dual tasks, there are two measurements to consider in detail. The first is the *dual-task deficit* which is a comparison of mean performance in the dual-task condition compared to the single-task condition. For perceptual tasks, this kind of dependency is often interpreted as a sign of limited-capacity, parallel processing in perception. The extreme version of this dependency is that only one stimulus can be processed at a time (serial processing).

A second measurement arises from requiring the same perceptual judgment on both stimuli. Consider the example of yes-no light detection: congruent stimuli are cases in which the light was presented to both sides or to neither side, and incongruent stimuli are cases in which the light was presented on one side and not the other. A *congruency effect* is a difference in performance between trials with congruent stimuli and those with incongruent stimuli. This effect is specific to the stimuli presented on a given trial rather than on mean performance. Such congruency effects are common in the selective attention literature where the intent is to measure failures of selective attention. Examples include the congruency effects found with the Simon paradigm (Simon & Rudell, 1969; Simon, 1990), the flanker paradigm (Eriksen & Eriksen, 1974; Eriksen & Hoffman, 1972; Yantis & Johnston, 1990), and the Stroop paradigm (Stroop, 1935; MacLeod, 1991). In all of these paradigms, one measures whether there is an effect of the relevant stimulus being congruent or incongruent with an irrelevant stimulus. In the divided attention literature, such congruency effects can be interpreted as either an error in selection (Yantis & Johnston, 1990) or a consequence of interactive processing such as crosstalk (e.g. Navon & Miller, 1987).

### Isolating perception

Our primary goal is to understand attention effects in perception. We have already described how the choice of this particular dual-task paradigm is motivated by the desire to study perception rather than memory, decision or response processes. But do the particulars of a given study allow such isolation? This will be addressed in follow-up experiments that test whether perception was isolated. To foreshadow part of our results, some effects, but not all, are shown to be specific to immediate stimulus processing such as perception and memory encoding.

## Background

### Previous Research on Dual-Task Deficits

Studies of dual tasks using accuracy measures that employ the same (or at least similar) perceptual judgment for both tasks began with Sperling and Melchner (1978) and Kinchla (1980) They introduced the idea of an attention operating characteristic (AOC) function that is discussed later in this article. To illustrate this approach, consider another early study using a pair of line-length tasks (Bonnel, Possamai & Schmitt, 1987). Observers were presented with two pairs of lines: one pair on the left and one pair on the right side of fixation. The lengths of the pairs of lines were either identical or slightly different. The observers were to judge whether the lengths of the pair were the same or different and give ratings of their confidence. The several blocked conditions included both single-task and dual-task conditions. The results were described by ROC functions and sensitivity was estimated by the *d’e* measure (for an introduction, see Macmillan and Creelman, 2005). In *d’* units, performance was 2.02±0.05 in the single-task condition and 1.28±0.05 in the dual-task condition. A clear cut dual-task deficit. Bonnel and colleagues interpreted the result as consistent with a fixed-capacity, parallel model (Shaw, 1980) which is discussed in detail below. Additional evidence for dual-task deficits have been observed for a variety of perceptual tasks including: pairs of luminance discriminations (Bonnel, Stein & Bertucci, 1992; Morrone, Denti & Spinelli, 2004); pairs of form discriminations (Lee, Koch & Braun, 1999; Ward, Duncan & Shapiro, 1996); pairs of color discriminations (Lee, et al., 1999; Morrone, et al., 2004), pairs of motion discriminations (Festman & Braun, 2010; Lee, et al., 1999); and, pairs of auditory identifications (Best, Gallun, Ihlefeld & Shinn-Cunningham, 2006; Gallun, Mason & Kidd, 2007).

Recently, White, Palmer and Boynton (2018) have examined accuracy dual tasks for masked words. For a semantic categorization task, they found large dual-task deficits that were consistent with an all-or-none serial model. Such a large deficit is unlike almost all of the other results cited above (but see the extreme example tasks in Sperling and Melchner (1978). We return to consider the unique results of this study in the discussion.

In contrast to studies that find dual-task deficits, there are two situations that often show no dual-task deficit. The first of these is when one must do two detection or detection-like tasks at the same time. For example, Bonnel et al. (1992) had observers detect luminance increments to the left or right of fixation and found no dual-task deficit. Similar results have been found for the detection of gratings at two different spatial locations and with two different spatial frequencies at the same location (Graham, Kramer & Haber, 1985); for orientation discriminations of Gabor patches in noise at two different locations (Han, Dosher, Lu, 2003; Liu, Dosher & Lu, 2009), and phase discriminations of Gabor patches in noise at two different locations (Han, et al., 2003). In all of these cases, the discrimination tasks were “detection-like” because the attribute being discriminated was quite different (e.g. horizontal vs. vertical orientation) so that discrimination thresholds were the same as detection thresholds (see studies of detection and discrimination investigating the concept of “labeled lines” (Watson & Robson, 1981).

The second situation with little or no dual-task deficit is when two properties of a single object are judged in a dual task (Duncan, 1984; Han, et al., 2003; Bonnel & Prinzmetal, 1998). For example, Bonnel and Prinzmetal (1998) presented letters to the left and right of fixation that varied in shape (F or T) and in color (green or blue). When the same letter was tested there was no dual-task deficit; but, when different letters were tested, there was a dual-task deficit. Thus, given the same pair of judgments, the occurrence of a dual-task deficit depended on whether the two tasks were on the same or different objects.

In summary, for a variety of perceptual tasks, one finds dual-task deficits when two judgments are required rather than one. However, there are at least two exceptions to this generalization. One is for detection or detection-like tasks and the other is for cases with two judgments of the same perceptual object. For these cases, there is little or no dual-task deficit.

### Previous Research on Congruency Effects in Dual Tasks

A congruency effect is the difference in performance on trials with congruent and incongruent stimuli. It is a dependency in the processing of the stimuli presented on a given trial and is predicted by theories with processing interactions between the two tasks such as crosstalk (Navon & Miller, 1987) or by theories with errors in selection (Yantis & Johnston, 1990; also called attentional slippage, Gaspelin, Ruthruff, Jung, 2014). Such a dependency is logically distinct from the dependency of mean performance measured by the dual-task deficit. One can also measure such a congruency effect in the single-task condition because it depends on only the presentation of both stimuli, not on making both responses. Such a congruency effect in the single-task condition is logically equivalent to the congruency effect measured for the flanker task. For a study that explicitly combines flanker and dual-task paradigms see Huber and Lehle (2007).

For speeded dual tasks, congruency effects were probably first measured by Hommel (1998, often called *backward crosstalk effects* in this literature) and have since received much study (e.g. Logan & Gordon, 2001). For the accuracy dual tasks with a common perceptual judgment, there have been only a few studies that have measured congruency effects. Perhaps the earliest was a set of auditory studies by Sorkin and colleagues (Gilliom & Sorkin, 1974; Sorkin, Pastore & Pohlmann, 1972; Sorkin, Pohlmann & Gilliom, 1973; and, Sorkin, Pohlmann & Woods, 1976) and a more recent auditory example is McCloy and Lee (2015). A good example from the visual literature is Bonnel, et al. (1992). They had observers view a display with two continuously illuminated lights to ether side of fixation. The intensity of each light was briefly (100 ms) incremented or decremented and the observers task was to press a corresponding key for an increment or a decrement. This modulation was simultaneous for the lights on the left and right side and whether it was an increment or a decrement was independent for the two sides. By this design, congruent stimuli were either both increments or both decrements. Incongruent stimuli were an increment on one side and a decrement on the other. Percent correct for the congruent condition was 97.5% and for the incongruent condition was 74.5% a difference of 23%.

There have been few other congruency effects reported for accuracy dual tasks with the same perceptual judgment in both tasks (e.g. Ernst, Palmer & Boynton, 2012). Part of the reason is that some dual-task studies use methods that minimize congruency. For example, the well-known study of Sperling and Melchner (1978) required the identification of single digits as the response. With 10 possible stimuli and corresponding responses, congruent trials make up 1/10 of the total while incongruent trials make up 9/10 of the total. Thus, measuring congruency effects becomes impractical. Another example is the use of different stimuli and judgments for the two tasks. In such cases, it may be ambiguous which stimuli are congruent (but see Long, 1975, 1977 for congruency effects between judgments of auditory frequency and of visual luminance). Such questions of correspondence are avoided by studying the case of interest here: dual tasks with a common judgment and simple two-choice responses. For that case, one can always test for congruency effects.

### Theories of Divided Attention

Consider now theories of divided attention that can account for both dual-task deficits and congruency effects. We focus on perception, but some role for memory, decision and response is a constant possibility. Consider four kinds of theory in order of increasing complexity.

#### Single resource theories

Many early theories of divided attention specify a single resource as the basis for the presence (or absence) of dual-task deficits and other effects of divided attention. Examples of such dependencies include: A processing bottleneck that allows only a single stimulus to be identified at a time (early selection theory, Broadbent, 1958); A single processing “channel” with limited capacity (single capacity theory, Kahneman, 1973); or, specific to vision, the orienting of perception to one location and not another (single spotlight metaphor, Posner, 1980).

By such theories, one expects dual-task deficits for pairs of tasks that compete for the limiting resource. In other words, a failure of independence (dependency) is governed by a resource such as whether an stimulus is selected, whether limited-capacity processing is devoted to the stimulus, or whether the spotlight shines upon it.

Within these theories, congruency effects are expected from errors in selection (Yantis & Johnston, 1991). If the wrong stimulus is processed on some fraction of the trials, it has consequences for incongruent stimuli but not congruent stimuli. Thus, these theories predict congruency effects when selection fails.

#### Interactive processing theories

A quite different theory of divided attention focuses on trial-by-trial dependencies due to processes interfering with one another. For example, processing simultaneous stimuli might result in *crosstalk*. This is when the processing of one stimulus is affected by the processing of the other stimulus. The term “crosstalk” is borrowed from the phenomena in electronic circuits in which the signal transmitted on one circuit creates an undesired effect on a nearby circuit. Examples of crosstalk in theories of perception and cognition include pooling models of perceptual crowding in which the stimulus representations are weighted averages that include contributions from nearby stimuli (Parkes, Lund, Angelucci, Solomon & Morgan, 2001), and response competition models in which competing response processes receive inputs from both stimuli (Eriksen & Schultz, 1979). Navon and Miller (1987) suggested that a variety of such interactive processes underlie both congruency effects and dual-task deficits. The idea is that the interaction results in a net loss of information that results in a decline of mean performance. A example of such an account is given by the all-or-none mixture model in Appendix B. Mordkoff and Yantis (1991) motivated such an interaction to take advantage of contingencies between stimuli. Indeed, Miller (1987) created the correlated flanker paradigm in which the flanker effects had to depend on contingencies. In sum, the interaction of processing during individual trials can predict both congruency effects and dual-task deficits. In the extreme, theories with widespread interactive processing can be thought of as having a single dependency because everything interacts.

This kind of theory turns the ideas of the single limited-capacity dependency on its head. In that case, dual-task deficits are due to limits on capacity which has a side effect of selection errors that cause congruency effects. For interactive processing theories, congruency effect are due to interactive processing which has a side effect in aggregation that cause dual-task deficits.

#### Theories of automaticity

Consider next the theory of automaticity in which there are two kinds of processes: controlled and automatic (Shiffrin & Schneider, 1977; Logan, 1988). Controlled processes allow the flexible processing of different sources of information at the price of the dependency of limited capacity. Automatic processes are inflexible stereotyped processes that operate on all stimuli and cannot be controlled. This makes them prone to interference from conflicting processes which is a different kind of dependency. In addition, automatic processes must be learned through experience while control processes can be assembled on demand. The result is that automatic processes do not contribute to dual-task deficits but do contribute to congruency effects. In contrast, control processes contribute to dual-task deficits but not to congruency effects. Thus, this theory predicts a tradeoff between incurring a dual-task deficit versus incurring a congruency effect. But, because more complex tasks can be a combination of both controlled and automatic processes, the details of the tradeoff depend upon an analysis of the processes required for a particular task.

Another way to look at automaticity theory is that it combines the dependency of a single resource theory with the dependency of an interactive processing theory. The controlled processes introduce one dependency due to limited resources while the automatic processes introduce a second dependency due to interactive processing. In summary, the theory of automatic and controlled processes is an example of a theory with two kinds of dependency that are differentially active for different tasks.

#### Multiple dependencies

A more general theory is that there are multiple limits on performance due to both processing capacity and interactive processing (e.g. Allport, Antonis & Reynolds, 1972; Allport, 1980; Pashler, 1989; 1991; 1998). One version of this idea specifies that these dependencies are specific to a module of processing. For example, there might be different dependencies for tasks that are limited by perceptual processes and tasks that are limited by response processes (Pashler, 1989). To focus on perception, one can allow for both limited-capacity processing of objects (e.g. object-based attention, Duncan, 1984) and spatially-local interactive processing of features (e.g. crowding, Parkes et al., 2001). Depending on the details of the display and the task, one, the other, or both dependencies might limit performance. And, of course, inadvertent bottlenecks or interactions in memory, decision or response processes might appear in what had been presumed to be a perceptual task. Hence, isolating perception and testing that isolation becomes even more important when the dependencies can be different for different modules. The challenge for these kinds of theories is to make them specific enough to have testable predictions.

#### Roadmap for more detailed theories

These general theories are pursued using more specific theories that are described following Experiments 1 and 2. In addition, formal definitions of the relevant quantitative models are in Appendix B.

## Summary of Goals

Our first goal is to test the predictions of single dependency theories of divided attention in perceptual tasks. We do this by measuring both dual-task deficits and congruency effects for two quite different perceptual judgments. Dual-task deficits and congruency effects must covary if they are to be accounted for by the same dependency in processing.

Our second goal is to provide a systematic analysis of the theories that might account for congruency effects. This goal grew as our study progressed and we found ways to distinguish among the theories in the literature.

## General Methods

### Two Perceptual Judgments

We investigated two perceptual judgments: detecting a simple visual pattern -- a Gabor patch -- or categorizing a single word. For *Gabor detection*, one had to detect a Gabor patch in visual noise. Did a particular display contain a horizontal Gabor patch in noise or did it contain just noise? For *word categorization*, one had to judge if a word belonged to a particular category. For example, one might have to judge whether a particular word represented an animal (e.g. “dog”) or a mode of transportation (e.g. “bus”). For both judgments, targeted and distractors were presented equally often and required a yes-no-like response in the form of a confidence rating.

### Procedure

The procedure is illustrated in Figure 1 which shows the stimulus sequence for the three conditions of our initial experiments. Consider first the *single-task condition* in the left column. A trial began by indicating the target category (e.g. “horizontal” for the Gabor detection and “animal” or other target categories for the word categorization). After a brief interval, the stimuli were presented in dynamic noise. The critical stimulus (e.g. Gabor patch) was brief (details below) within a noise display of 1 s duration. There were noise displays on both sides for this condition, a block of trials always had the relevant stimulus on one side. Thus, the observer’s task was to judge just one side (hence *single-task condition*). After a short interval to avoid masking, there is a prompt to respond. Each observer was assigned a cue color (red or blue) and was to respond according to the stimulus on the side with the cued color. For the example illustrated in the figure, the blue cue is on the left and accordingly an observer with the blue cue instruction is to respond to the stimulus on the left. This color cue is used so that the displays have no purely visual differences between the left and right sides. For the single-task condition, this response cue was always on the same side for the entire block. The trial ended with a response in the form of a confidence rating and tone feedback was given for errors.

**Figure 1.**
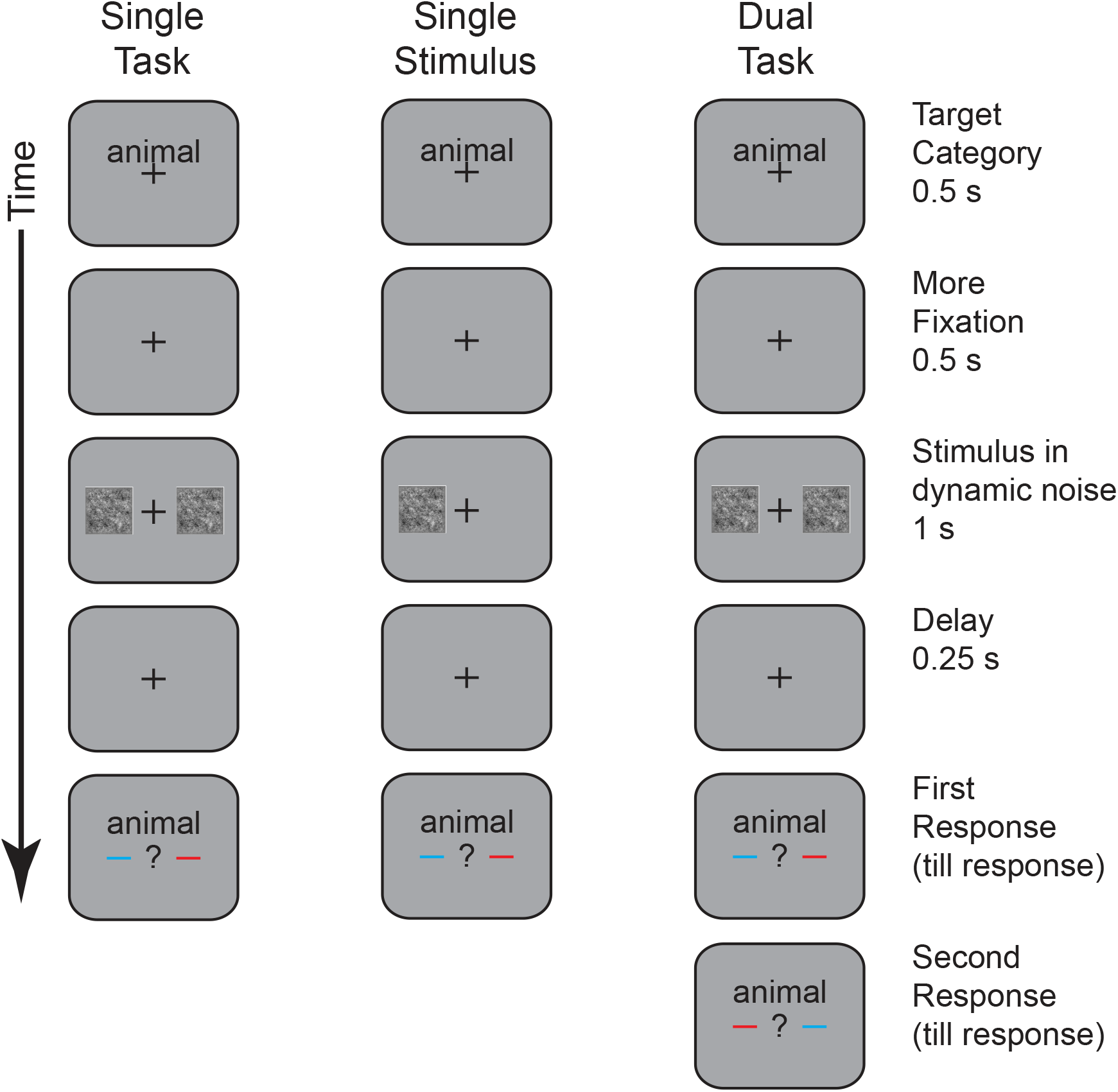
An illustration of the general procedure. The stimulus sequence is shown for the three main conditions: single task, single stimulus, and dual task. All conditions begin with a fixation display that includes the target category. After a brief delay, the first stimulus is displayed within a 1 s movie of dynamic 1/f noise. Then after another brief delay, the observer is prompted for a response using a post cue that specifies the relevant side of display for this response. In the single-task condition, the relevant side is blocked and the observer is informed at the beginning of the block. In the single-stimulus condition, everything is the same except the there is no noise movie on the irrelevant side. In the dual-task condition, both sides are relevant for every trial of a block. The display sequence is identical to the single-task condition, but with both sides tested in sequence. In this example, the blue cues indicate the relevant side.

Next consider the *dual-task condition* shown in the rightmost column. The displays were identical up to the response cue. A block of the dual-task condition differed from a block of the single-task condition in having tests of both the left and the right side. In some of the experiments, two responses were required, one after the other. The side that was prompted first is unpredictable to prevent the preparation of a response while the stimuli are presented. In sum, the single-task and dual-task conditions had identical displays but differed in that the observer must perform one perceptual task (e.g. left side detection) versus two perceptual tasks (e.g. separate detection judgments of the two sides).

The third *single-stimulus condition* is shown in the middle column of the figure. The task in this condition was the same as with the single-task condition: Judge an entire block of trials with the relevant displays on either the left or right side. The distinctive feature was to remove the irrelevant display. This allows one to test if the presence of a second irrelevant display has any effect on performance. This single-stimulus control is a check for failures of selective attention that might reduce mean performance.

The critical stimuli (Gabors or words) were presented briefly with temporal uncertainty during the relatively long dynamic noise display. Specifically, the stimulus contrast was modulated by a Gaussian temporal waveform that had its peak during the noise display and a standard deviation of 0.05 s. The peak was restricted to not occur in the first or last 0.2 s of the display. The effective duration of this stimulus was about 0.1 s in the 1 s display of noise. The time that the target appeared for the two tasks was the same to prevent observers from switching the attended side after seeing one target. This synchrony of target presentation was the only way in which the physical stimuli for the two tasks are dependent on one another.

The spatial structure of the display is shown in Figure 2. The two noise patches were 6 by 6° to either side of a 0.5° fixation cross. They were each centered at an eccentricity of 4° which resulted in an 2° space between them. Overall, the two noise patches filled the middle 14° of a video monitor that had a viewable width of about 32°. An example Gabor patch is shown in the right side with a contrast of 80% which is much higher than used in all but the last experiment. It was presented with spatial and temporal uncertainty in the noise display. For example, the Gabor stimuli had an Gaussian envelope with a standard deviation of 0.5°. This made them effectively about 1° in size. The stimuli were excluded near the edge of the display (< 0.5°) to prevent clipping the and the noise was attenuated to prevent sharp edges. As a result, the stimuli appeared anywhere in a region of 5 by 5° (25 square degrees).

**Figure 2.**
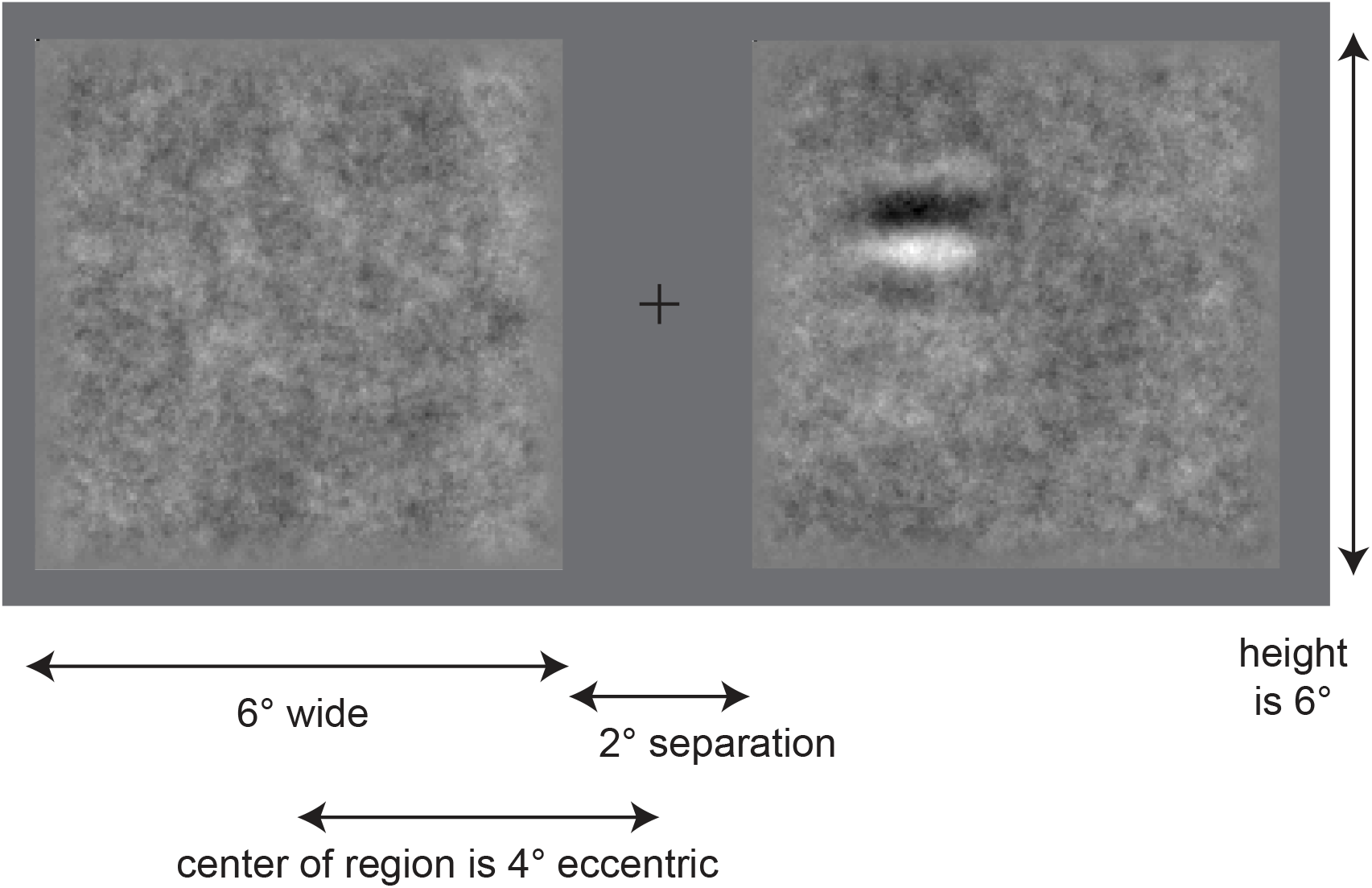
An illustration of a single frame of the stimulus display. Two examples of the 1/f noise are shown on each side of fixation. The display on the right includes a high contrast Gabor patch (80%). The figure also specifies the dimensions of each display element.

Stimuli were presented within a dynamic 1/f noise display. Individual pixels had luminance values that were initially independently sampled from a Gaussian distribution and were then filtered in space and time so that each dimension had 1/f noise. The luminance values of each pixel had a distribution with a mean at zero contrast and a standard deviation of 12% contrast. New noise frames were presented at a rate of 30 Hz (every 4th refresh of our 120 Hz display). In summary, the contrast for component frequencies varies inversely with the frequency. Thus the noise has relatively more low frequency content than white noise. This kind of noise is useful because it equates the “power” per octave which is more relevant to human vision than equating the power per degree as in white noise. Thus, 1/f noise is an effective kind of noise for stimuli with a wide range of spatial and temporal scales.

Observers were instructed to maintain fixation during the stimulus displays. To enforce this, we measured eye position during all trials using an eye movement monitor. Trials with changes in eye position of more than 1° horizontal or vertical were aborted. Overall, there were less than 3% aborts. More details about eye position are described with the apparatus.

The three main conditions (dual-task, single-task, single-stimulus) are blocked. In addition, the side for the single-task and single-stimulus conditions is blocked. This yields 5 kinds of blocks: dual-task; left-single-task; right-single-task; left-single-stimulus; right-single-stimulus. To equate the number of trials in the primary conditions, there were 2 dual-task blocks along with 1 each of the 4 other kinds of blocks.

### Analysis of Responses

Observers responded with one of four key presses that indicated likely-no, guess-no, guess-yes, or likely-yes. These ratings were used to form a receiver operating characteristic (ROC) function and performance was summarized by the percent area under the ROC (*A*_*ROC*_). For reasonable assumptions, this *A*_*ROC*_ measure is equivalent to the percent correct measured by a forced choice paradigm (Green & Swets, 1966). To estimate *A*_*ROC*_ we used the simple “connect the dots” method to avoid making distributional assumptions (also called the trapezoid method, Macmillan & Creelman, 2005).

### Details of Gabor Detection

Several details of the procedure differed for the two judgments. For Gabor detection, the stimuli were either a noise image alone or a noise images with a single horizontal Gabor patch. Observers judged the presence or absence of the Gabor patch. Thus there was a consistent mapping between stimuli and response which is believed to encourage automatic processing of the stimulus (e.g. Shiffrin & Schneider, 1977). The grating component of the horizontal Gabor patches were in sin phase with a spatial frequency of 1 c/d. The Gaussian envelope had a standard deviation of 0.5° and was truncated to a maximum size four times the Gaussian standard deviation (4 × 0.5 = 2° across). The contrast of the Gabor was adjusted for each observer with values from 18-35%.

### Details of Word Categorization

A goal of using word categorization was to be sure that observers were reading the words for meaning and not making a decision based on the presence of a few letters. One way to help insure such reading is to change the target category from trial to trial so that there is no consistent mapping between details of the stimuli and the response. This is called variable mapping and was used for our word judgments. Thus, at the beginning of every trial, the initial display of the target category informed the observer what is the target category for that trial. Such variable mapping between stimuli and response is believed to encourage controlled processing of the stimulus (e.g. Shiffrin & Schneider, 1977).

We used six word categories (Scharff et al. 2011a) with 8 words each that are given in Table 1. They were presented in the fixed size Courier font at 36 point which under our viewing conditions yielded characters that fit within a rectangle of 0.85 by 1.41°. Thus, the longest word (5 characters) was at most 4.25° wide and 1.41° high. The characters were displayed as dark strokes on the middle gray background. The target contrast of the words was adjusted for each observer resulted in values of 18-35%.

**Table 1.**
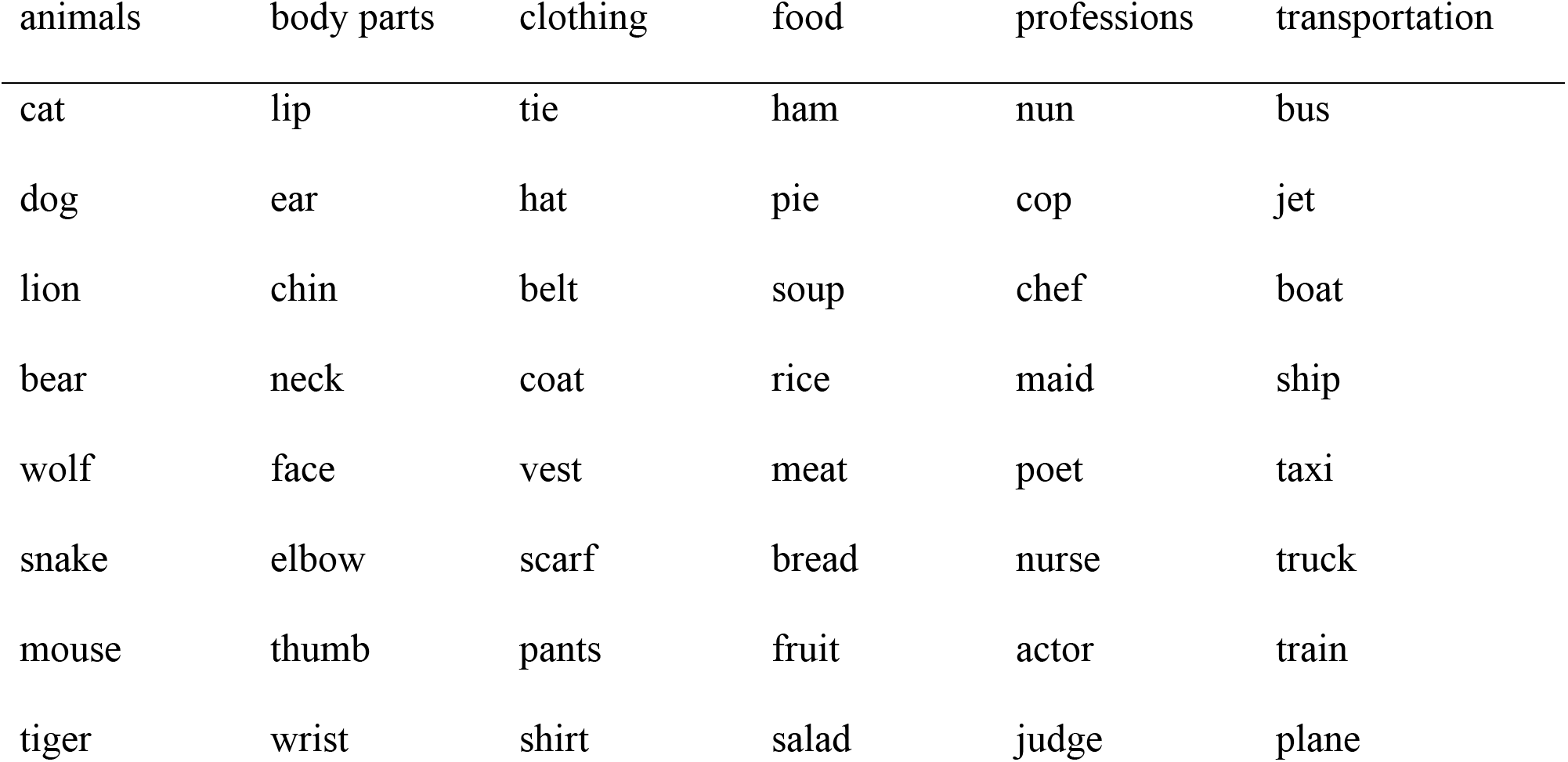
Words used in the Experiment

Words used in the Experiment
| animals | body parts | clothing | food | professions | transportation |
| --- | --- | --- | --- | --- | --- |
| cat | lip | tie | ham | nun | bus |
| dog | ear | hat | pie | cop | jet |
| lion | chin | belt | soup | chef | boat |
| bear | neck | coat | rice | maid | ship |
| wolf | face | vest | meat | poet | taxi |
| snake | elbow | scarf | bread | nurse | truck |
| mouse | thumb | pants | fruit | actor | train |
| tiger | wrist | shirt | salad | judge | plane |

A further detail of the word experiments involved how distractors were chosen. Depending on the details, it is possible to introduce contingencies between the stimuli that might contribute to congruency effects. For example, Mordkoff (1996; see also Mordkoff and Yantis, 1991) have shown that some versions of the flanker paradigm introduced contingencies between the flanker and target stimuli that contribute to the flanker effect. This has also been shown in the Stroop paradigm, Melara & Algom, 2003) and makes contact with the larger study of contingency learning (e.g. Lin & MacLeod, 2017).

For our word experiments, there were six categories and on any given trial, one category defined the targets and the other five defined the distractors. A target was presented on half of the trials and this probability was independent on the two sides. For the distractors, we chose to pick a single distractor category for a given trial to present for both tasks on both sides. Thus, if there was a distractor on both sides, it was always from the same category. This introduced a contingency between distractor categories. If one knew the distractor category on one side, then one would know it on the other.

In retrospect, we could have chosen distractors in simpler ways. For example, each trial could have been a choice between a single target category and a single distractor category to make the contingencies relatively simple. But for current purposes, the contingencies in the word experiments are probably irrelevant. This is because there were little congruency effect for words. Thus, the absence of a substantial congruency effect makes moot the argument about it being due to inadvertent contingencies.

In summary, the two judgments differed in several ways: the detection of a particular Gabor patch with consistent mapping of target and distractor, versus the categorization of many possible words with variable mapping of target and distractor.

### Aspects of the Procedure motivated by our Imaging Experiments

Two aspects of this procedure were intended to increase the size of a fmri signal examined in a separate study (White, Runeson, Palmer, Ernst & Boynton, 2017). The spatial extent of the noise display was relatively large (6×6°) and nearly all of it is relevant to the judgment due to the spatial uncertainty of the target. The duration of the noise display was relatively long (1 s) and nearly all of it is relevant due to the temporal uncertainty of the target. In summary, the large and long noise displays provided a potent signal for our related fmri study.

### Observers

In most experiments there were 6 observers. Many were in multiple experiments and over the series of experiments there were a total of 11 observers. Some were unpaid volunteers and others were paid $20/hr. All had normal or corrected-to-normal vision.

### Apparatus

The stimuli were displayed on a flat-screen CRT monitor (19” ViewSonic PF790) controlled by a Power Mac G4 (Dual 1.0 GHz) using Mac OS X 10.6.8. The experiment was displayed at a resolution of 832 × 624 pixels, a viewing distance of 60 cm (25.5 pixel/degree at screen center), and a refresh rate of 120 Hz. The monitor had a peak luminance of 119 cd/m², and a black level of 4.1 cd/m², mostly due to room illumination. Stimuli were displayed using Psychophysics Toolbox 3.0.11 for Matlab R2012a (Brainard, 1997). A chin rest with an adjustable chair ensured a fixed distance to the display.

On all trials, eye position was recorded using an EyeLink II, 2.11 with 250 Hz sampling (SR Research, ON). The EyeLink II is a head-mounted binocular video system and was controlled by software using the EyeLink Developers Kit for the Mac 1.11.1 and the EyeLink Toolbox 3.0.11 (Cornelissen, Peters, & Palmer, 2002). The position of the right eye was recorded for all trials, and trials were included in the analysis only if fixation was confirmed. When fixation failed, five consecutive high frequency tones were sounded and the trial was aborted. The percentage of aborted trials for each observer in each experiment ranged from 0.5% to 5.2% with an overall mean including all experiments of 2.4 ± 0.2%. Thus the observers maintained fixation on almost all trials and none of the analyses included trials with blinks or saccades to the stimuli.

### Final Comment on Methods for Dual-Task Experiments

We have employed a refinement intended to help isolate the role of perception in divided attention effects. Specifically, response cues were used to indicate which response to make rather than using a predictable order of responses. Using such a cue prevents an unintended prioritization of one response over the other. It also can prevent effects due to preparing the first response while still perceiving the other stimulus. In a previous study (Ernst, et al., 2012), we found in pilot work that there was an order effect when the responses were in a fixed order but not when using this unpredictable response cue. In addition, fixed order cues were used in at least some of the studies that argue against independent processing for simple detection tasks (Pastukhov, Fischer, Braun, 2009).

## Experiment 1

### Methods

The first experiment measured divided attention effects for detecting Gabor patches. As just described, there were 3 blocked conditions: single task, single stimulus, and dual task. In addition, the data from the dual-task condition were broken down by the first or second response. Thus, for our initial analysis we considered 4 main conditions: single task with two stimuli, single task with a single stimulus, first-response dual task, and second-response dual task. After practice, there were 6 observers who participated for 5 hours resulting in 640 trials in each of the 4 main conditions for each observer.

### Results

Performance in the 4 main conditions are shown in the left panel of Figure 3. Performance is measured in terms of the percent area under the ROC function. As described in the methods, this measure can be thought of as an estimate of the unbiased percent correct. There was little difference between the 4 conditions. The difference between the single-task conditions with two or one stimuli was near zero (0.1±1.0%, t(5)= 0.12, p>.1). The difference between the first and second responses for the dual-task condition was also near zero, opposite of the expected direction and is not reliable by a two-tailed test (−0.8±0.4%, t(5)=1.92, p>.1). Combining across these control conditions, the dual-task deficit was 1.3 ± 0.9% which was marginally reliable (t(5)=1.62, p<.1). In summary, the controls revealed no reliable effects and there was little or no dual-task deficit for Gabor detection. This was what was expected based on prior studies of dual-tasks using detection judgments (e.g. Bonnel, et al., 1992).

**Figure 3.**
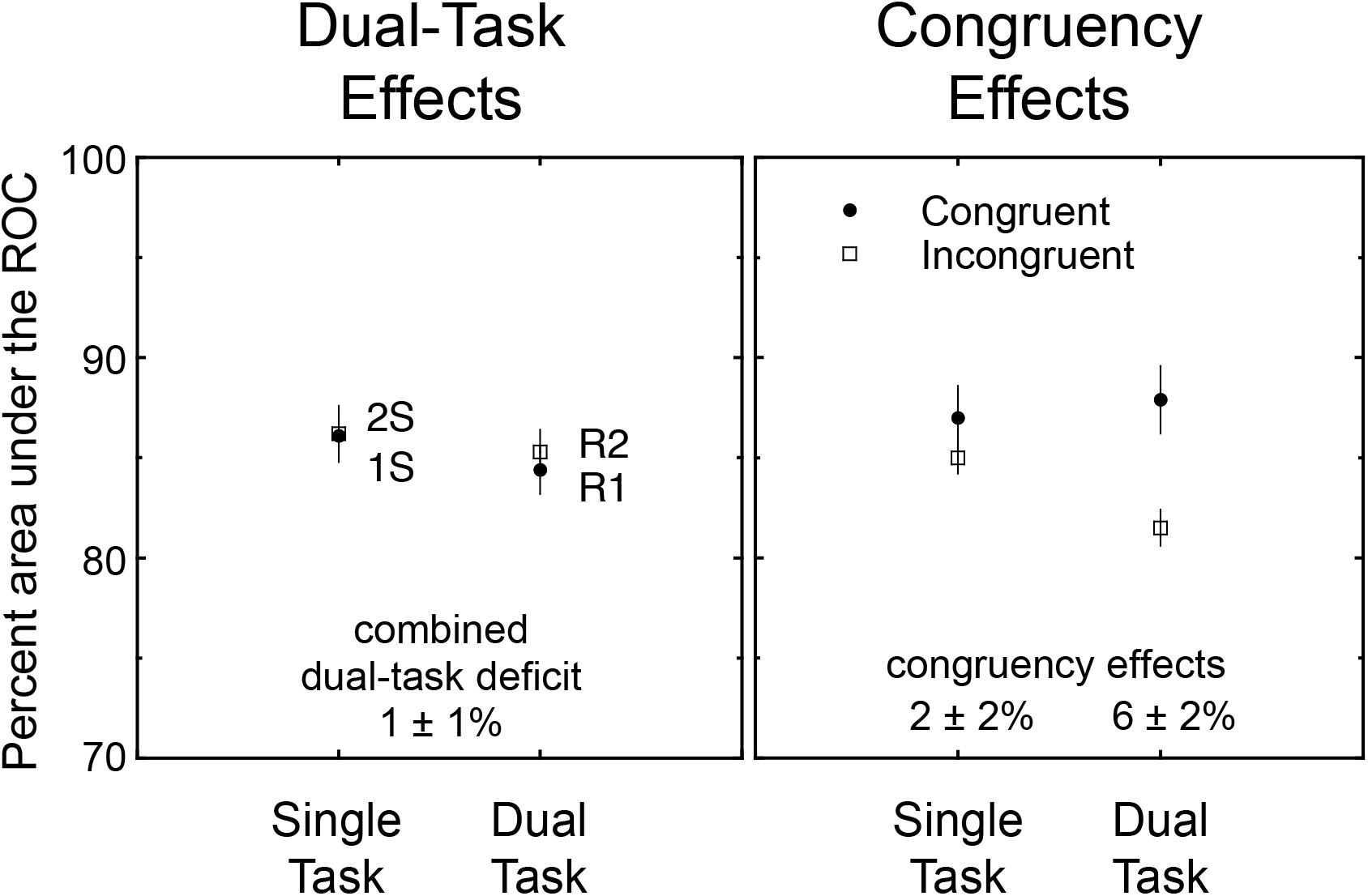
Results of Experiment 1. In the left panel, percent area under the ROC is shown for four conditions: single-task (2S), single-stimulus (1S), and the dual-task condition shown separately for the first (R1) and second response (R2). The area under the ROC can be thought of as an unbiased measure of percent correct. The dual-task deficit (difference between the single- and dual-task conditions) was relatively small and not reliable. In the right panel, percent area under the ROC is plotted for two main conditions: the single-task and dual-task conditions combined over response order. These conditions are further broken down by whether the trial had congruent or incongruent stimuli. The congruency effect (difference between congruent and incongruent) was reliable for the dual-task condition but not the single-task condition.

Now consider congruency effects. The stimuli in the two tasks are congruent if they require the same response. The effect of congruency is shown in the right panel of Figure 3 for the single-task condition (two stimuli) and for both responses of the dual-task condition. For the single-task condition, the congruency effect was small and not reliably different than zero (2.1 ± 1.4%, t(5)=1.45, p>.1). In contrast, the dual-task congruency effect was larger and was reliable 6.4 ± 1.6% (t(5)=3.89, p<.01). Thus, even though there was no reliable dual-task deficit in the dual-task condition, a judgment on one side was affected by the stimulus on the other side and thus yields a congruency effect.

## Experiment 2

### Methods

In the second experiment, divided attention effects were measured for categorizing words. As with Experiment 1, there were 4 main conditions: single task with two stimuli, single task with a single stimulus, first-response dual task, and second-response dual task. After practice, six observers participated for 5 hours resulting in 640 trials in each of the 4 main conditions for each observer.

### Results

In the left panel of Figure 4, performance in terms of percent area under the ROC is shown for the 4 main conditions. As with Experiment 1, there was no effect of the presence of a second stimulus or of response order. The difference between the single-task conditions with two or one stimuli was 0.8±0.7% (t(5)= 1.15, p>.1). The difference between the first and second responses for the dual-task condition was −0.2±1.3%, (t(5)= 0.17, p>.1). Combining across these control conditions, there was a reliable dual-task deficit of 5.4 ± 1.1% (t(5)=4.77, p<.005). In summary, unlike the Gabor detection task, word categorization had a dual-task deficit of about 5%.

**Figure 4.**
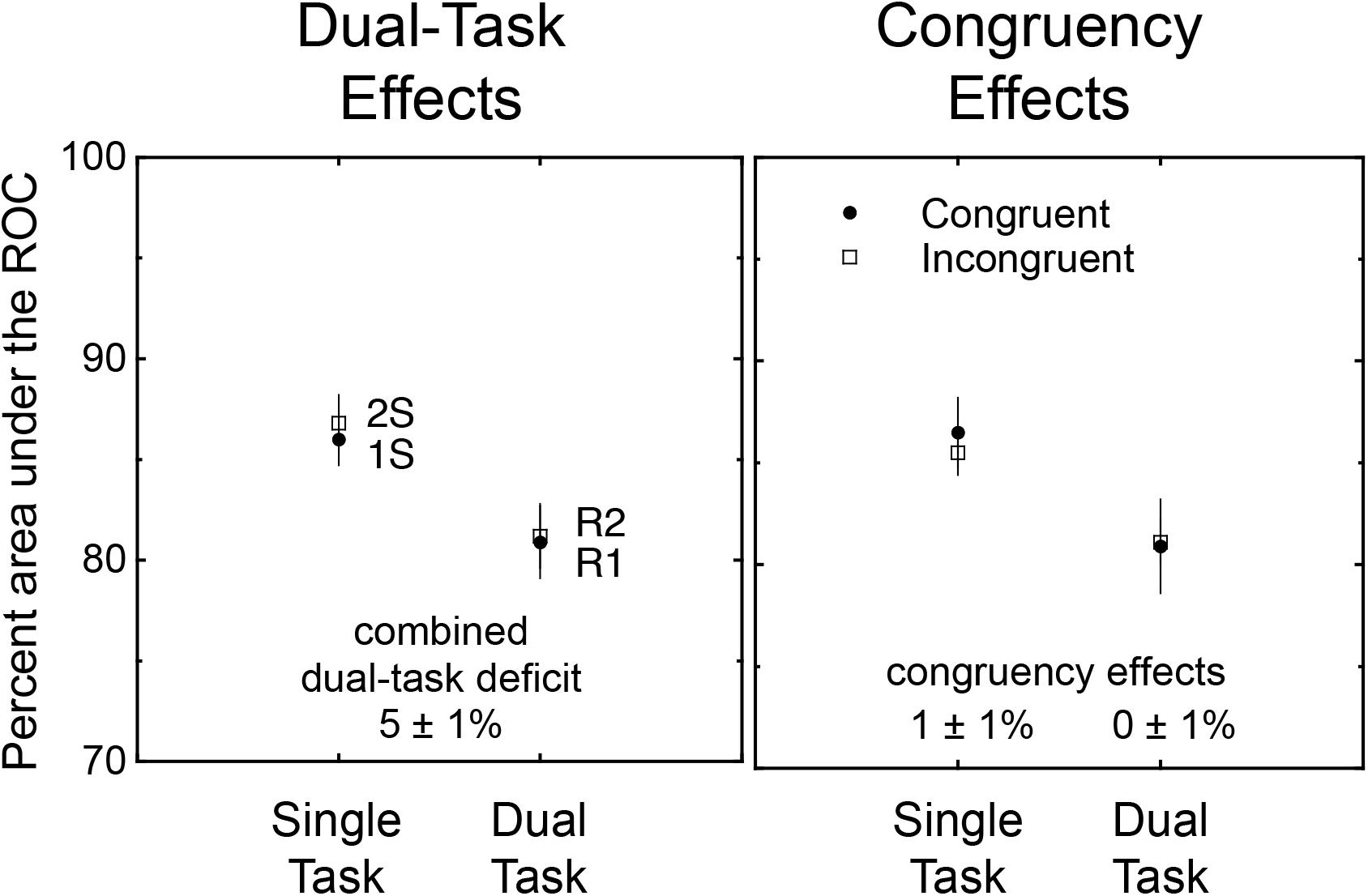
Results of Experiment 2. In the left panel, percent area under the ROC is shown for four conditions: single-task (2S), single-stimulus (1S), and the dual-task condition shown separately for the first (R1) and second response (R2). The dual-task deficit was relatively large and reliable. The congruency effects of Experiment 2. In the right panel, percent area under the ROC is plotted for two main conditions: the single-task and the dual-task conditions combined over response order. These conditions are further broken down by whether the trial had congruent or incongruent stimuli. The congruency effect was not reliable for either of the conditions.

The congruency effects for this experiment are shown in right panel of Figure 4. On the left side are the results for the single-task condition (two stimuli) and on the right side are the results for both responses of the dual-task condition. Unlike Gabor detection, neither effect was reliable. The single-task congruency effect was 0.9 ± 1.1% (t(5)=0.86, p>.1). The dual-task congruency effect was −0.2 ± 1.7% (t(5)=0.13, p>.1). Thus, for word categorization there were little or no congruency effects even for the dual-task condition.

## Models of Dual-Task Deficits

### Alternative Models

We next introduce several models of divided attention that have been used to construct theories of both visual search and dual tasks. In the following section, these models are elaborated to include an account of congruency effects. The early development of these models was in the context of response time (Townsend, 1990; Townsend & Ashby, 1983). Here, we focus on their consequences for accuracy experiments (Scharff et al., 2011a).

#### Common assumptions

We emphasize a handful of relatively specific models that provide landmarks among the large set of possible models. These landmark models make specific predictions and can thus be easily tested. In addition, two more general models are described to guide further discussion. They all share the following assumptions.

Consider judgments that require distinct responses for displays with target stimuli and distractor stimuli. We assume that the evidence that a particular stimulus is a target corresponds to a single random variable. Unless otherwise specified, these random variables are identically distributed and independent of one another. One can think of these random variables as corresponding to the aggregate output of all relevant perceptual processes. The decision process uses these random variables to determine a response. Following signal detection theory (Green & Swets, 1966), the appropriate decision rule can be relatively simple. For example, for a yes-no judgment, the independent decision rule is based on comparing a criterion value to the value of the random variables on that trial. If there is more evidence than the criterion, then respond “yes”; if not, respond “no”.

#### Unlimited-capacity, parallel model

In this simplest of models, the stimuli are processed independently and a decision is made by the independent decision rule. It predicts no dual-task deficit. This model is described in detail in Appendix B.

#### Fixed-capacity, parallel model

In this model, the processing of each stimulus is assumed to be parallel but there is a dependency in the quality of information obtained about each stimulus. Specifically, a constant amount of information is obtained per unit time (or per display). This information theory definition can be understood in terms of an equivalent process model such as the sample size model (Shaw, 1980). In the sample size model, one takes *n* independent samples from the relevant stimuli in the display and the precision of perception depends on the number of samples taken from a particular stimulus. When only a single stimulus is relevant, then all samples can be taken from that stimulus. When two stimuli are relevant, the samples must be distributed over the two stimuli. Given equal relevance, there would be *n/2* samples per stimulus. Consider now the random variable that results from taking the mean of the samples from a given stimulus. The standard deviation of the mean of the samples varies as the square root of the number of samples. Thus, the standard deviation for stimuli in the dual-task condition is a square root of two larger than for stimuli in the single-task condition. Given further assumptions about the distribution (e.g. Gaussian), one can calculate the dual-task deficit expected given any level of performance in the single-task condition.

#### Limited-capacity, parallel model

This model is a generalization of the two models just described. Assume there is some degree of dependency in the processing of multiple stimuli similar to the sample size model. The degree of dependency can range from none (unlimited capacity) to fixed capacity and to even more limited. To quantify the model for the dual-task case, we use a generalization of the sample size model that is described in Appendix B. In brief, the parameter *k* specifies the degree to which capacity is limited. A value of *k*=*0* specifies an unlimited-capacity model, and a value of *k*=*1* specifies the fixed-capacity model. Values in between specify a continuous range of models with intermediate limits on capacity. In addition, values of *k*>*1* specify a dependence even greater than expected from the fixed-capacity model. This model can predict any magnitude for the dual-task deficit.

#### All-or-none serial model

This model differs in that only one stimulus is processed on a given trial. But an observer can switch what stimulus is processed from trial to trial. Thus, one ends up with a mixture of trials in which the relevant stimulus was attended and other trials in which the relevant stimulus was not attended (Davis, Shikano, Perteron & Michel, 2003; Dosher, Han, & Lu, 2004; White et al., 2018). The idea of this all-or-none serial model has been implemented in models that go by several names including the all-or-none mixture model (type 1 in Shaw 1980) and the all-or-none switching model (Bonnel & Prinzmetal, 1998). This model predicts relatively large magnitude dual-task deficits and also predicts deficits that persist even for very discriminable stimuli.

#### Standard serial model

This model is a generalization of the all-or-none serial model and is closer to the serial models discussed in the response time literature. Versions of this kind of model were called the *type 2 mixture model* by Shaw (1980) and the *probabilistic serial search model* by Dosher, et al. (2004). For these models, there is some fraction of trials when both stimuli are processed. When that fraction is zero (only one stimulus processed), it reduces to the all-or-none serial model. When that fraction is one (both stimuli always processed), the model becomes equivalent to the unlimited-capacity, parallel model. As with the limited-capacity, parallel model, this model can predict any magnitude of dual-task deficits. But it does differ in how dual-task deficits vary with discriminability. Unlike the parallel model, high discriminability cannot overcome a serial process that does not complete both stimuli on some fraction of the trials.

#### Summary

Three of the models described provide specific predictions about the magnitude of the dual-task deficits: the unlimited-capacity, parallel model; the fixed-capacity, parallel model; and, the all-or-none serial model. The more general limited-capacity, parallel and standard serial model do not make such specific predictions and must be distinguished by more subtle properties such as response correlations, and how the magnitude of the dual-task deficit varies with discriminability.

### The Magnitude of the Dual-Task Deficit

Consider next the magnitude of the dual-task deficits predicted by the models. The first step in this analysis is to replot the results using an attention operating characteristic (AOC, Sperling & Melchner, 1978; Sperling & Dosher, 1986). The idea of the AOC is shown in Figure 5. The two panels on the left illustrate the basic results of a dual-task experiment: Accuracy is plotted for the single-task and dual-task conditions broken down by whether the response was to the left side or the right side. In this example, both sides have a single-task accuracy of 85% and a dual-task accuracy of 77%. In the right most panel is the AOC for the same data. An AOC is a parametric plot of the performance for one task against the performance of a second task. Specifically, the single-task accuracy is shown on the axes and the dual-task accuracy is shown in the interior of the plot. In other words, the x,y value of this interior point are the accuracy in the dual-task condition for the left- and right-sides, respectively. The 4 values in the simple plot on the left are rearranged for the AOC into 2 values along the axes and 2 values to define the interior point.

**Figure 5.**
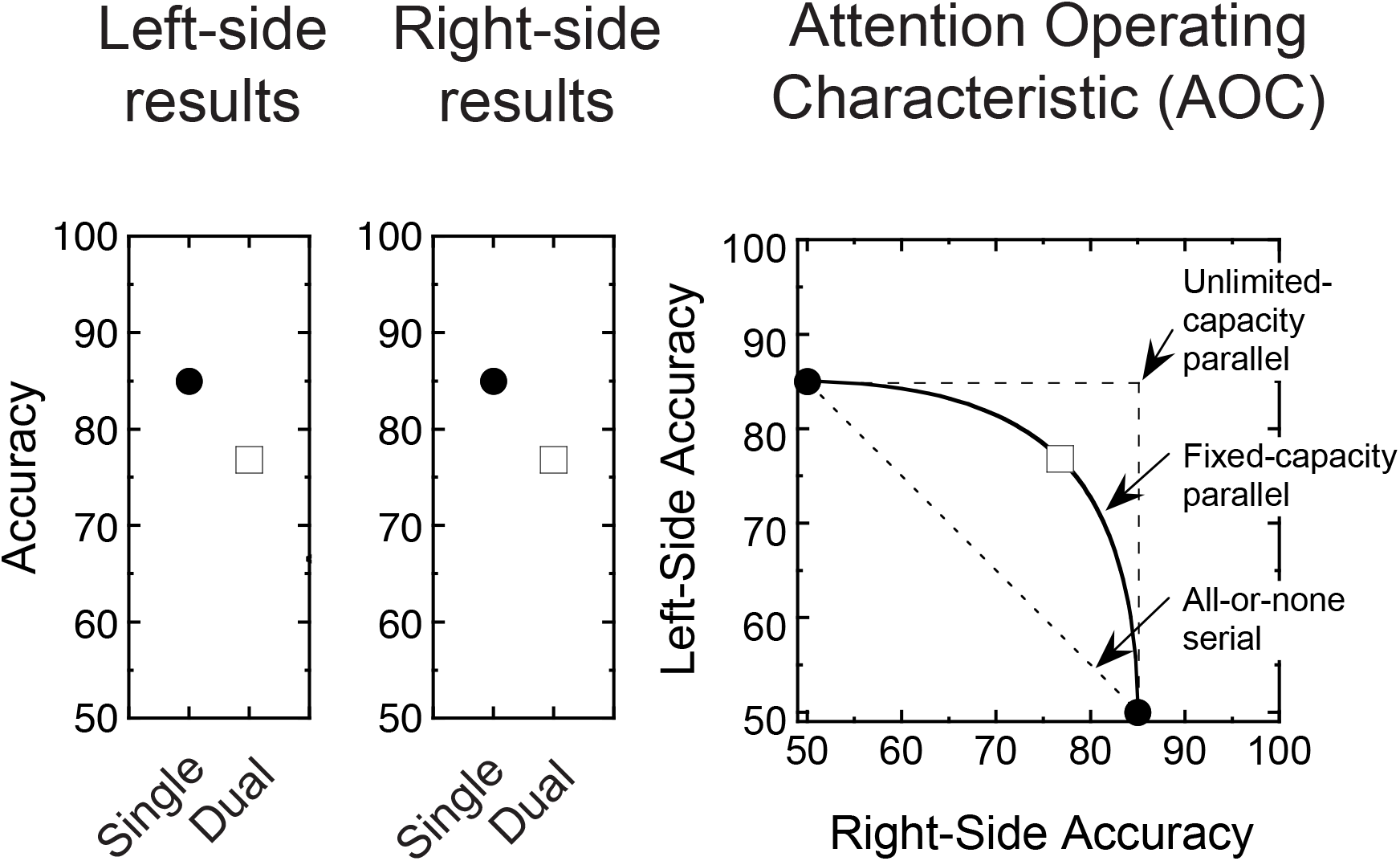
Illustration of the construction of an attention operating characteristic. The two panels on the left show simple plots of single- and dual-task conditions for the left and right side. The panel on the right shows the same data replotted as an attention operating characteristic. This is a parametric plot with left-sided accuracy plotted against right-sided accuracy. The single-task performance is plotted on the axes and the dual-task performance is plotted in the interior of the plot. The predictions for three models are shown: unlimited-capacity, parallel, fixed-capacity parallel, and all-or-none serial. The example data matches the prediction of the fixed-capacity, parallel model.

The advantage of the AOC is that it is easy to illustrate the predictions of the three landmark models described earlier in this article. (a) An unlimited-capacity, parallel model predicts no dual-task deficit. Consequently, it predicts dual-task performance at the *independence point* defined by the two dashed lines that trace out the values of the single-task conditions. (b) A fixed-capacity, parallel model with independent Gaussian noise predicts a result with performance somewhere along the solid curve shown in the figure. Where along the curve depends on the allocation of capacity between the two tasks. In this example, equal allocation results in the performance shown by the open square symbol. (c) An all-or-none serial model predicts performance somewhere along the dotted line between the two single-task results. In sum, by plotting the results for the two sides against one another, one can reveal the predictions of the landmark models for dual-task deficits.

The AOCs for the first two experiments are shown in the two panels of Figure 6. The Gabor detection experiment is shown in the left panel and the word categorization experiment is shown in the right panel. For Gabor detection, the performance for dual tasks fell just inside of the independence point and well above the curve for the fixed-capacity, parallel model. Thus there is little if any capacity limitation.

**Figure 6.**
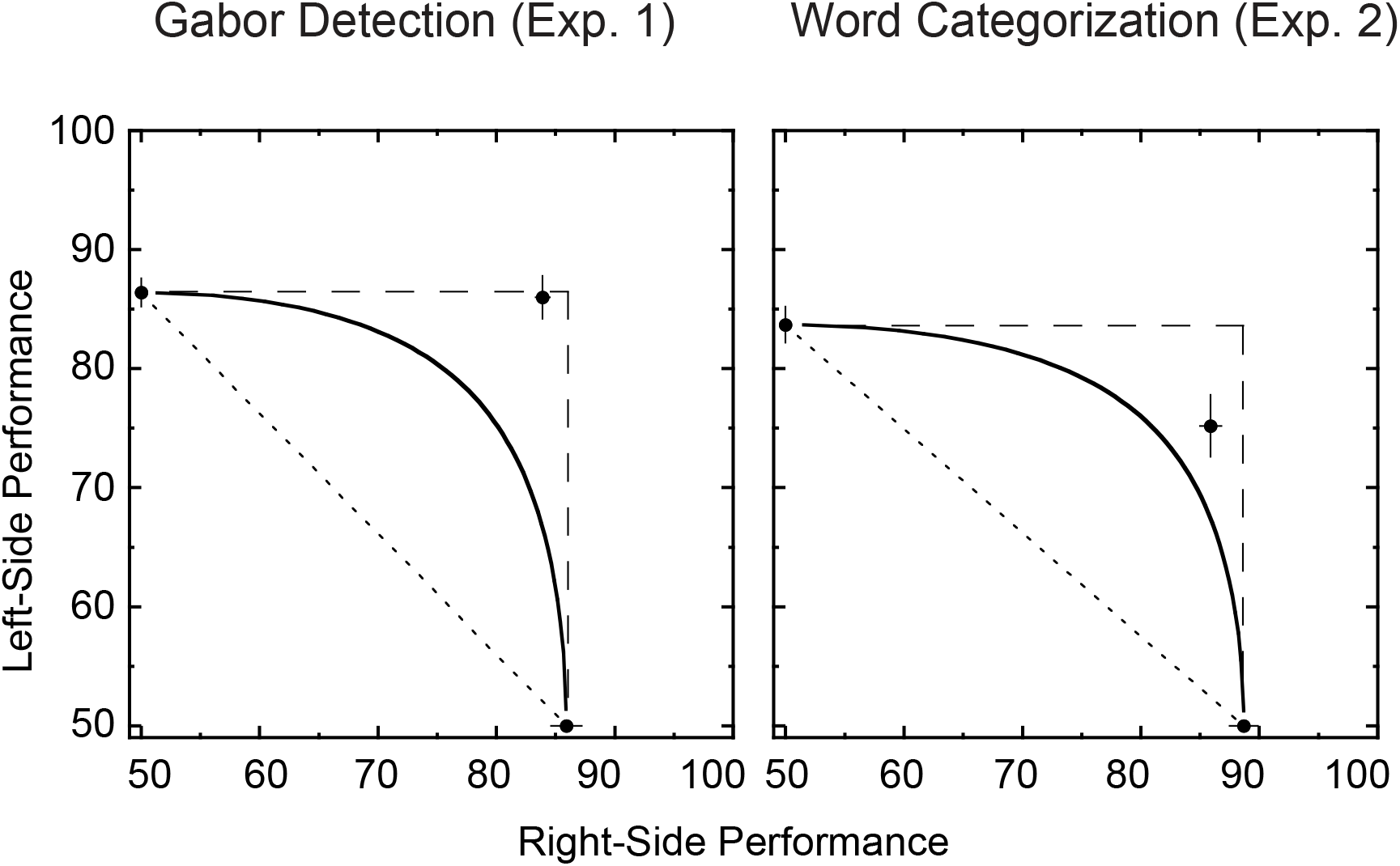
AOCs for Experiments 1 and 2. The top panel shows the AOC for Gabor detection from Experiment 1 and the bottom panel shows the AOC for word categorization from Experiment 2. For Gabor detection, the dual-task point was near the independence point consistent with an unlimited-capacity model. For word categorization, the dual-task point was well away from the independence point but above the fixed-capacity curve. This is consistent with limited-capacity processing.

For word categorization, the performance for dual tasks fell roughly halfway between the independence point and the curve for the fixed-capacity, parallel model. For this task, capacity is limited but the limitation is less than that predicted by a fixed-capacity, parallel model and not nearly as limited as the prediction of the all-or-none serial model.

In summary, plotting the results on AOCs puts the magnitude of the capacity limitations into a meaningful context. For Gabor detection, the limitation is relatively small. For word categorization, the limitation is about half that predicted by a fixed-capacity, parallel model.

### Response Correlations

In dual-task conditions, one can measure two responses on a single trial. This allows one to ask whether these responses are correlated in any way. Sperling and Melchner (1978) have pointed out that parallel and serial models make different predictions about such correlations. In particular, serial models typically predict negative correlations between a correct response on one task and a correct response on the other task. Parallel models in themselves predict no correlation. But, any common noise source for the two tasks would introduce a positive correlation. Thus, a negative correlation stands out as a signature of a serial process. Quantitative predictions of response correlations are given in Appendix B for selected parallel models.

The aggregate response contingencies for Gabor detection are given in Table 2. The rows specify if the left-side task was correct and the columns specify if the right-side task was correct. For example, if one is correct on both tasks, that trial is counted in the lower right cell. The value in each cell is the proportion of trials with that particular response pair. This table aggregates across all observers and includes 3823 response pairs. The correlation for each individual can be estimated from their contingency table. The mean correlation was .044 ± .015 for the six observers (t(5)= 2.9, p<.05). A small but reliable positive correlation. One can also consider the correlation broken down by the kind of trial. For signal-signal trials it was .05±.03. For signal-noise trials, it was −.01±.01. And for noise-noise trials, it was .17±.07. Thus trials with two distractors (and no targets) had the largest positive correlation. This pattern among the correlations was also found for the color tasks in White et al. (2018).

**Table 2.** Response Contingencies in Experiment 1 (Gabor detection)

| Left-side response | Right-side response |  |
| --- | --- | --- |
|  | Error | Correct |
| Error | .043 | .136 |
| Correct | .158 | .664 |

The aggregate response contingency table for word categorization (Experiment 2) is shown in Table 3 with 3741 response pairs. The mean correlation estimated from individual observers was .020 ± .005 (t(5)=4.0, p<.05). We also analyzed the correlations for different types of trials and found only a hint of negative correlation (signal-signal, −.02±.02; signal-noise, .10±.03; and noise-noise, −.05±.08). In sum, both experiments have small positive correlations between the two responses. Thus, for this task, there was no sign of the negative correlation expected from a serial model. For dual-task experiments that find the negative correlation, see Sperling and Melcher (1978) or White, et al. (2018).

**Table 3.** Response Contingencies in Experiment 2 (word categorization)

| Left-side response | Right-side response |  |
| --- | --- | --- |
|  | Error | Correct |
| Error | .060 | .235 |
| Correct | .129 | .575 |

## Theories of Congruency Effects

### Three Distinctions among Theories

There are a variety of theories for congruency effects in part because they arise in many domains including both selective and divided attention. We organize them in terms of three distinctions (Moore & Palmer, 2018). Two of these distinctions are illustrated in Figure 7. The first, and most conceptually difficult distinction, is between selection errors and interactive processing. With *selection errors*, congruency effects arise due to the selection of information from irrelevant stimuli: The observer meant to attend one stimulus, but by error attended the other stimulus (Yantis & Johnston, 1990). In contrast, with *interactive processing* the representation of one stimulus is influenced by other stimuli (Navon & Miller, 1987). As a result, even if one selects perfectly, an irrelevant stimulus influences the judgment of the relevant stimulus. Of course, these accounts are not mutually exclusive. Both kinds of dependency might contribute to a full account of congruency effects. For example, selection might modulate interactive processing to yield a hybrid account. In the figure, this distinction is shown by the separate columns.

**Figure 7.**
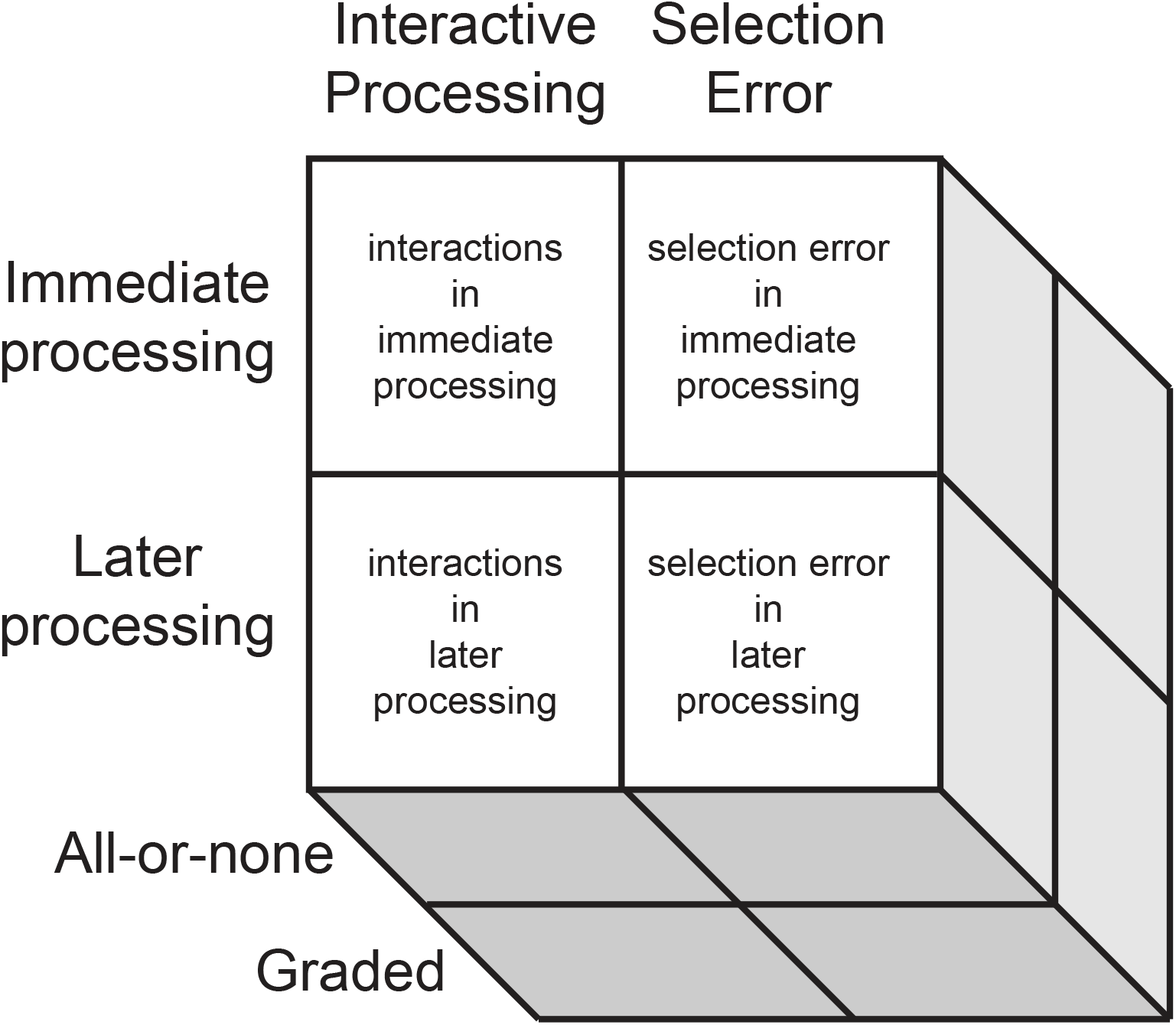
An illustration of three distinctions among alternative theories of congruency effects. The columns distinguish theories that depend on interactive processing or errors in selection. The rows distinguish theories with loci in the immediate processing following a stimulus rather than later processing. And the cells in depth distinguish theories with the critical process on a given trial being all-or-none or graded.

The second distinction is between cases were the congruency effect arises from immediate processing or later processing. Immediate processing refers to any process that immediately follows the stimulus and cannot be delayed to occur later in time. The primary examples of delayable processes are memory retrieval, decision and response processes. While immediate processing obviously includes much of perception, it also includes elements of memory (memory encoding) and possibly elements of decision (perceptual categorization) that must immediately follow the presentation of an stimulus. For example, memory encoding must occur while information from the stimulus is available. This immediate-later distinction is similar to the distinction between early selection (Broadbent, 1958) and late selection (Duncan, 1980). This distinction is also not mutually exclusive. Congruency effects might be due to dependencies in both immediate and later processing. In the figure, this distinction is shown by the separate rows.

The third distinction is whether the relevant process (selection or interaction) is graded or all-or-none. Examples of graded processes include pooling in perceptual crowding, weighting in decision and priming in response preparation. Examples of all-or-none processes include perceptual crowding with substitution of one percept for another, memory interference by substitution of one memory trace for another, and response competition by substitution of one response for another. “All-or-none” refers to the action on a given trial. Such all-or-none processes still predict gradual effects on performance when aggregated over trials. This third distinction is shown in depth in the figure. We present two models in Appendix B that capture the difference between graded and all-or-none processes.

### Four Theories of Congruency Effects

#### Selection error in immediate processing

This theory accounts for congruency effects as due to errors in a selection process active during or immediately following stimulus presentation. This theory evolved from early selection theory (Broadbent, 1958). In particular, Yantis and Johnston (1990) proposed that congruency effects in their version of the flanker task were due to errors in selection. This idea was pursued by Lachter, Forster and Ruthruff (2004) who called it slippage theory. For a recent review see Gaspelin, Ruthruff and Jung (2014).

#### Selection error in later processing

This theory also accounts for congruency effects by errors in selection. But now selection occurs well after stimulus presentation such as at time of memory retrieval or decision. This theory evolved from late selection theory (Duncan, 1980). The idea of slippage that was applied to early selection can be applied equally well to selection in late processing. An early example of this kind of theory is to make a decision on the basis of a weighted integration of evidence from the relevant and irrelevant stimuli (Kinchla & Collyer, 1974; Kinchla, Chen & Evert, 1995). This idea has been pursued in cueing paradigms (weighted linear integration, Shimozaki, Eckstein, & Abbey, 2003), spatial filtering (contrast gain and weighted integration, Palmer & Moore, 2009) and visual search (overlapping channels, Busey & Palmer, 2008).

#### Interactions in immediate processing

By this theory, selection is good enough that it doesn’t cause congruency effects. Instead congruency effects arise from some kind of interactive process such as crosstalk (Navon & Miller, 1987). A specific version for immediate processing can be found in theories of crowding that pose pooling between nearby stimuli (Parkes, et al., 2001) or by substitution among nearby stimuli (Ester, Klee & Awh, 2014).

#### Interactions in later processes

From the earliest papers on the flanker paradigm, congruency effects have been accounted for by response competition (Eriksen & Eriksen, 1974; Eriksen & Schultz, 1979). By this theory, response processes compete and congruent conditions allow more response preparation than incongruent conditions. This general idea has been elaborated in the speeded dual-task literature (e.g. Hommel, 1998; Logan & Gordon, 2001). Mordkoff and Yantis (1991) also elaborated this hypothesis with their interactive race model. In addition, one can easily imagine how an interactive processes in memory might cause congruency effects (interference theory, Oberauer & Lin, 2017). In sum, there are many theories about how interactions in later processing might cause congruency effects.

### Critical Properties

Each of these distinctions can be evaluated using appropriate experimental manipulations. Selection error and interactive processing can be distinguished by manipulating cues or instructions that affect selection while using otherwise identical stimuli, responses and tasks. In our study, two stimuli were always presented, but observers were cued to perform either a single task involving one of the stimuli or a dual task involving both. The stimuli were unchanged. Consider first how this manipulation affects selection. In a single-task condition, one can select information from the relevant stimulus from the beginning of a trial. In the dual-task condition, one must process both stimuli until the response cue allows the selection of information from the relevant stimulus. Now consider how this manipulation affects interactive processing. Because the stimuli and judgment are unchanged any “pure” interactive process are unaffected by the cue. Thus, any congruency effect that is due to an interactive process will be the same for single- and dual-task conditions. Finally, consider an example hybrid theory. The “early” selection allowed by the single task can protect the stimulus representation from interactions that would occur in the dual task. Thus, for this hybrid theory both selection and interaction play a role. In summary, cue manipulations allow one to distinguish pure interactive processing from either pure selection or a hybrid theory. In the current article, the effects of single- vs dual-task cues in Experiment 1 rule out a pure interactive processing account for congruency effects in Gabor detection.

Immediate and later processing can be distinguished by comparing simultaneous and sequential presentation of the stimuli (Shiffrin & Gardner, 1972). If immediate processing mediates the congruency effect, then sequential presentation should eliminate the congruency effect. If later processing mediates the congruency effect, then sequential presentation should result in the same congruency effect as simultaneous presentation. And, finally, if both loci are involved, then one expects that sequential presentation would reduce the congruency effect but not eliminate it. These predictions are tested in Experiments 3 and 4.

Graded and all-or-none processing can be distinguished by comparing low contrast conditions to conditions with high contrast that result in essentially perfect performance for the single-task condition (Palmer & Moore, 2009). All-or-none models are unusual in predicting the persistence of congruency effects at high contrast. Such a test is pursued in Experiment 7.

## Experiment 3

In the this experiment, we tested whether the divided attention effects found with Gabor detection are due to immediate processing such as perception or memory encoding or later processing such as decision that is not tied to the presence of the stimulus. The approach was to compare a simultaneous display of two stimuli to sequential displays in which the stimuli were displayed in sequence. In the visual search literature, this comparison has been used to test if the dependency between the stimuli in specific to immediate processing (Shiffrin & Gardner, 1972; Scharff, et al., 2011a, b). The strategy has been used less often with dual tasks (Duncan, 1980).

### Methods

Experiment 3 combined the dual-task and single-task conditions from Experiment 1 with two new conditions as shown in Figure 8. The four conditions are in separate columns. The leftmost column is the single-task condition which is unchanged from Experiment 1. The second column is the simultaneous dual-task condition. This was also unchanged except for measuring only one of the two possible responses.

**Figure 8.**
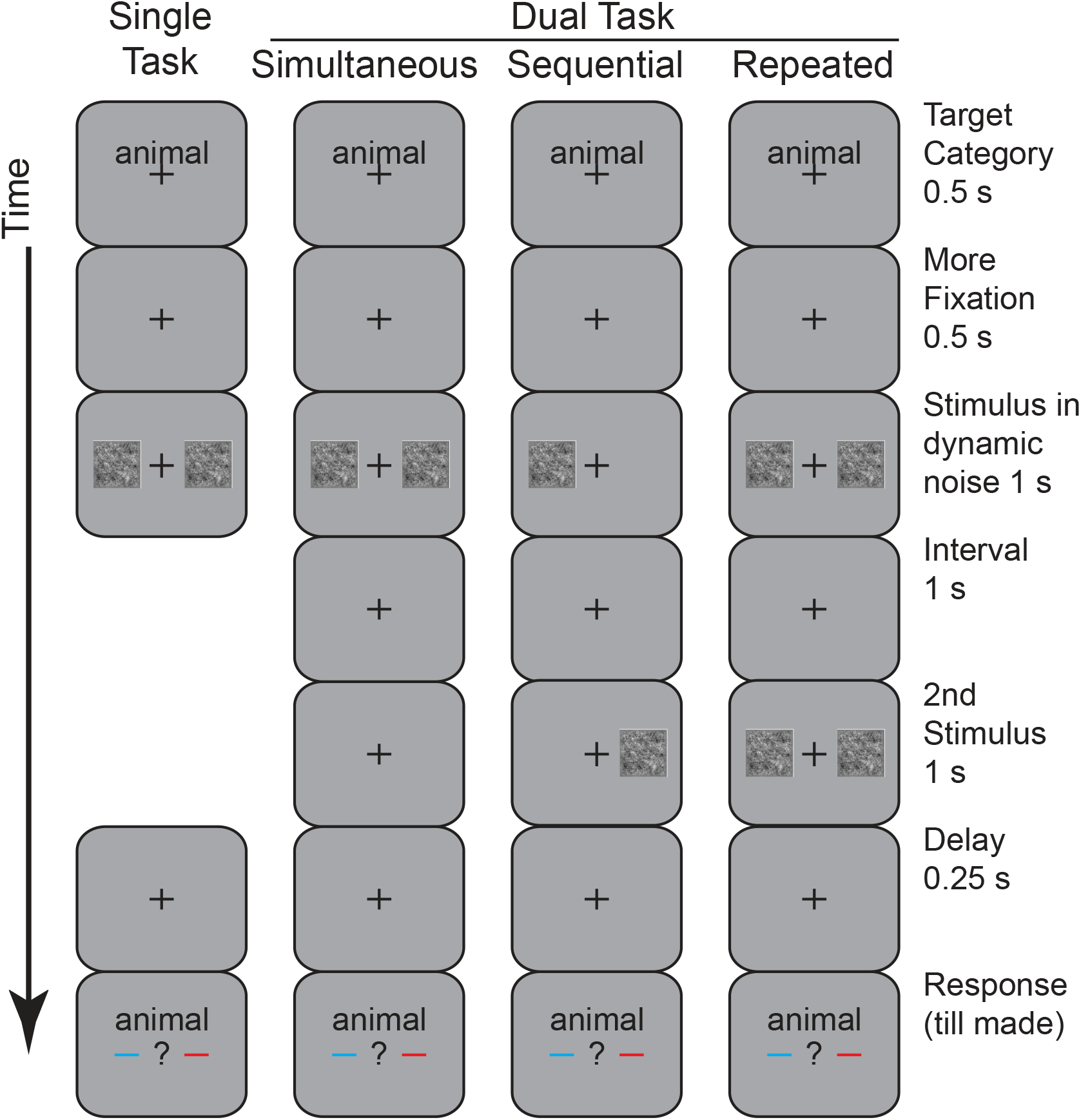
Illustration of the procedure of Experiments 3 and 4. The stimulus sequence is shown for the four conditions: single task, simultaneous dual task, sequential dual task, and repeated dual task. All conditions have the same initial and ending displays as the previous experiments. The single-task and simultaneous dual-task conditions are the same as the previous experiments. In the sequential-dual-task condition, the stimuli for the left and right sides are presented in separate displays with an intervening interval of 1 s. In the repeated-dual-task condition, the display for the simultaneous dual-task condition is repeated in a second display.

The third condition is called the *sequential dual-task condition*. The new feature is that the critical stimulus display is split into a pair of displays shown in sequence. In this example, the left-side stimulus was shown first and the right-side stimulus was shown second. This order of left or right displays was blocked. The duration of the individual displays was unchanged (1 s). The interval from the end of the first display to the beginning of the second display was 1 s. Such a long interval is likely to provide sufficient time to shift selection from display to display (Ward, Duncan & Shapiro, 1996). The logic of adding this condition is that if the dependency between the tasks is specific to immediate processing, then it will disappear for this sequential condition. On the other hand, if the dependency in not due to the processing of the immediate stimulus such as having to make the final decision, then it should be unchanged between the simultaneous and sequential displays.

The fourth condition is called the *repeated dual-task condition*. It also had two sequential displays. But these displays repeat the entire simultaneous display rather than split them apart. This purpose of adding this condition is to provide a comparison for the expected size of the dual-task deficit. For a class of fixed-capacity models, the difference in performance between dual and single tasks (dual-task deficit) is predicted to match the difference between the repeated and simultaneous dual-task condition (Scharff, et al., 2011a; 2013). To get an intuition, assume a serial model that can process only a single stimulus in the brief displays of this experiment. Then the dual-task deficit arises because in the dual-task condition only one of the two stimuli in the dual-task condition can be processed while in the single-task condition it is sufficient to process the one relevant stimulus. Similarly, the repeated effect arises because only one of the two stimuli are processed in the simultaneous dual-task condition while both of the stimuli are processed in the repeated dual-task condition. In other words, the repeated condition gives an observer a second chance at the second stimuli which makes it as good as the single-task condition.

In summary, this experiment combined the dual-task paradigm with a comparison of simultaneous and sequential displays. There were 4 main conditions: single task, simultaneous dual task, sequential dual task and repeated dual task. After practice, six observers participated for 7 hours resulting in 672 trials in each of the 4 main conditions for each observer.

### Predictions

The predictions for three different hypotheses are shown in Figure 9. For each panel, performance is shown as a function of the four conditions: single task, simultaneous dual task, sequential dual task, and repeated dual task. To emphasize particular comparisons, the shaded area marks the baseline performance defined by the simultaneous-dual-task condition. For example, the dual-task deficit can be visualized as the amount by which the single-task condition rises above the shaded area defined by the simultaneous-dual-task condition.

**Figure 9.**
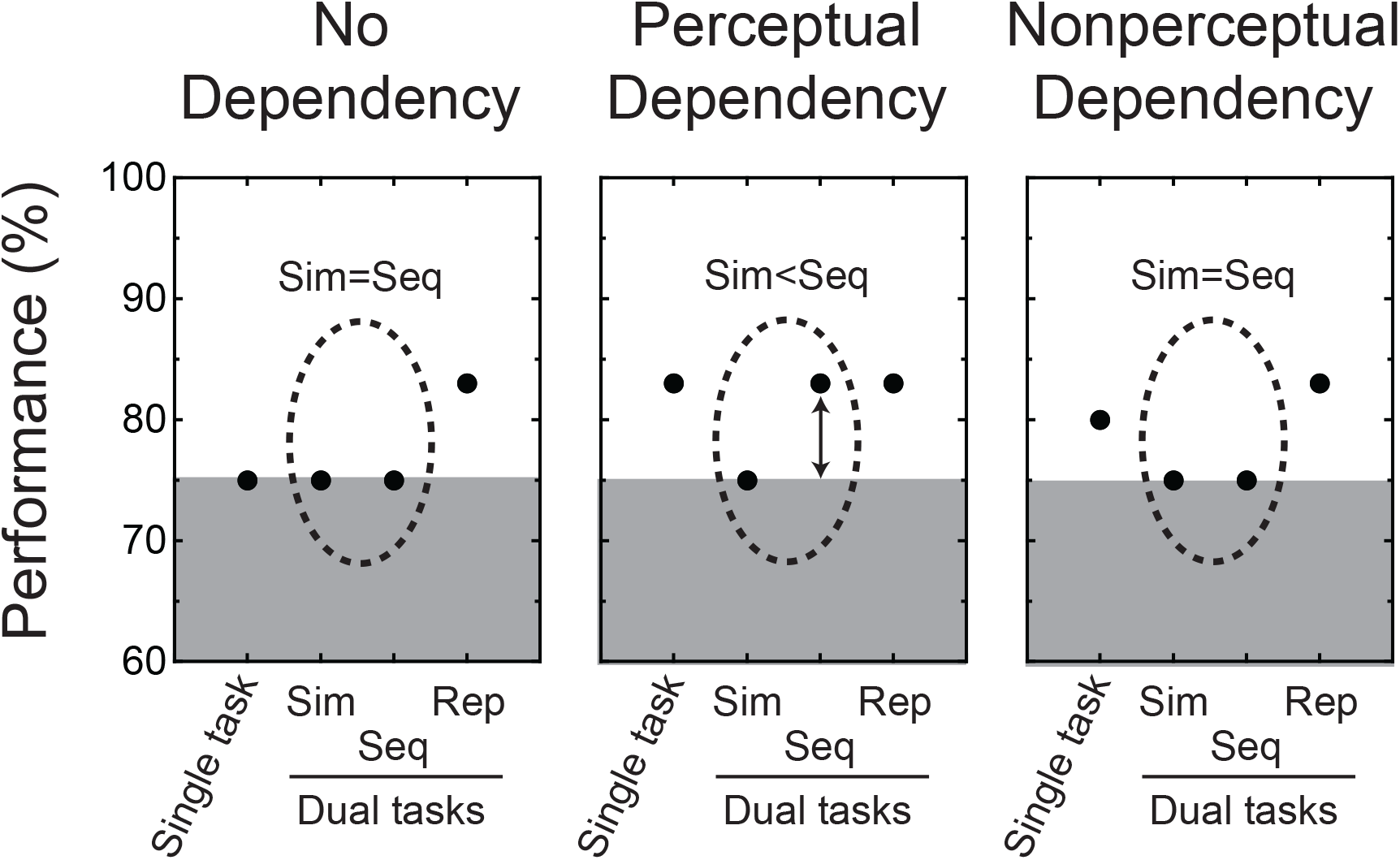
Predictions for Experiments 3 and 4. The three panels illustrate three different patterns of performance. In each, performance is shown for four conditions: single task, simultaneous dual task (sim), sequential dual task (seq), and repeated dual task (rep). The leftmost panel shows the case of no dependency between the two tasks (unlimited capacity). The single, simultaneous dual task and sequential dual task all yield the same performance with better performance for the repeated dual task. The middle panel shows the case of a immediate processing dependency between the two tasks (fixed capacity). The single, sequential dual task and repeated dual task all yield the same performance with worse performance for the simultaneous dual task. The rightmost panel shows the case of a dependency between the two tasks in later processing. The simultaneous dual task and sequential dual task yield the same performance with better (but not necessarily equal) performance for the single task and repeated dual task. For purposes of inferring a dependency in immediate processing, the key comparison is between simultaneous and sequential conditions which are circled in the illustration.

The leftmost panel shows the prediction for the case where there is no dependency between the two tasks. For this case, performance is the same for the single-task, simultaneous-dual-task and sequential-dual-task condition. The new feature for this case is that even without a dependency one expects an advantage for the repeated-dual-task condition. The second view of the stimuli improves performance. If this doesn’t happen, it undermines the logic of the simultaneous-sequential comparison. The task must allow an additional display to improve performance or the sequential-dual-task condition is not a test of capacity limits.

The middle panel shows the case with a perceptual dependency. Now the dual-task deficit is due to competition among immediate processes for the two stimuli. The leftmost two conditions show a dual-task deficit (single task > simultaneous dual task) as expected for such a dependency. The key prediction is the comparison highlighted by the dashed ellipse. Performance in the sequential dual task should be better than in the simultaneous dual task. This *sequential effect* is the hallmark of a dependency in immediate processing. Finally, the simplest version of this dependency predicts the sequential effect is the same size as the repeated effect. Namely, the performance in the sequential dual task equals that in the repeated dual task.

The rightmost panel shows the case with a nonperceptual dependency. Now the reason for a dual-task deficit is not immediate processing. Instead it is due to a dependency in memory, decision or some other process that is not tied to directly processing the stimulus display. Once again the leftmost two conditions show a dual-task deficit (single task > simultaneous dual task) and the key prediction is highlighted by the dashed ellipse. Performance in the sequential dual task should be the same as in the simultaneous dual task. Even though the stimuli are presented in sequence, the dependency in memory, decision or elsewhere remains. Unlike the perceptual dependency, there is no obvious reason to think this dependency should match the advantage of the repeated-dual-task condition. Accordingly, the performance in the sequential-dual-task condition is shown as not matching that in the repeated-dual-task condition.

### Results

In the left panel of Figure 10, performance in terms of percent area under the ROC is shown for the 4 conditions: single task, simultaneous dual task, sequential dual task and repeated dual task. Performance was similar for the single-task, simultaneous dual task, and sequential dual task. In addition, performance was reliably better for repeated dual task. The shading emphasizes the simultaneous dual-task condition as a baseline upon which to compare the other conditions. Using this baseline, consider three comparisons that are highlighted by giving the values at the bottom of the figure: the dual-task deficit (single-simultaneous) was 2.3±0.5%, (t(5)= 5.00, p<.01); the sequential effect (sequential-simultaneous) was −1.6±0.8%, (t(5)= 2.02, p<.1, two tailed); and, the repeated effect (repeated-simultaneous) was 7.2±0.5%, (t(5)= 15.5, p<.001). Thus, there was a substantial repeated effect combined with small dual-task deficits and sequential effects that go in opposite directions. The repeated effect confirmed that an additional display can improve performance, and the small size of the other two effects were consistent with a little or no dependency.

**Figure 10.**
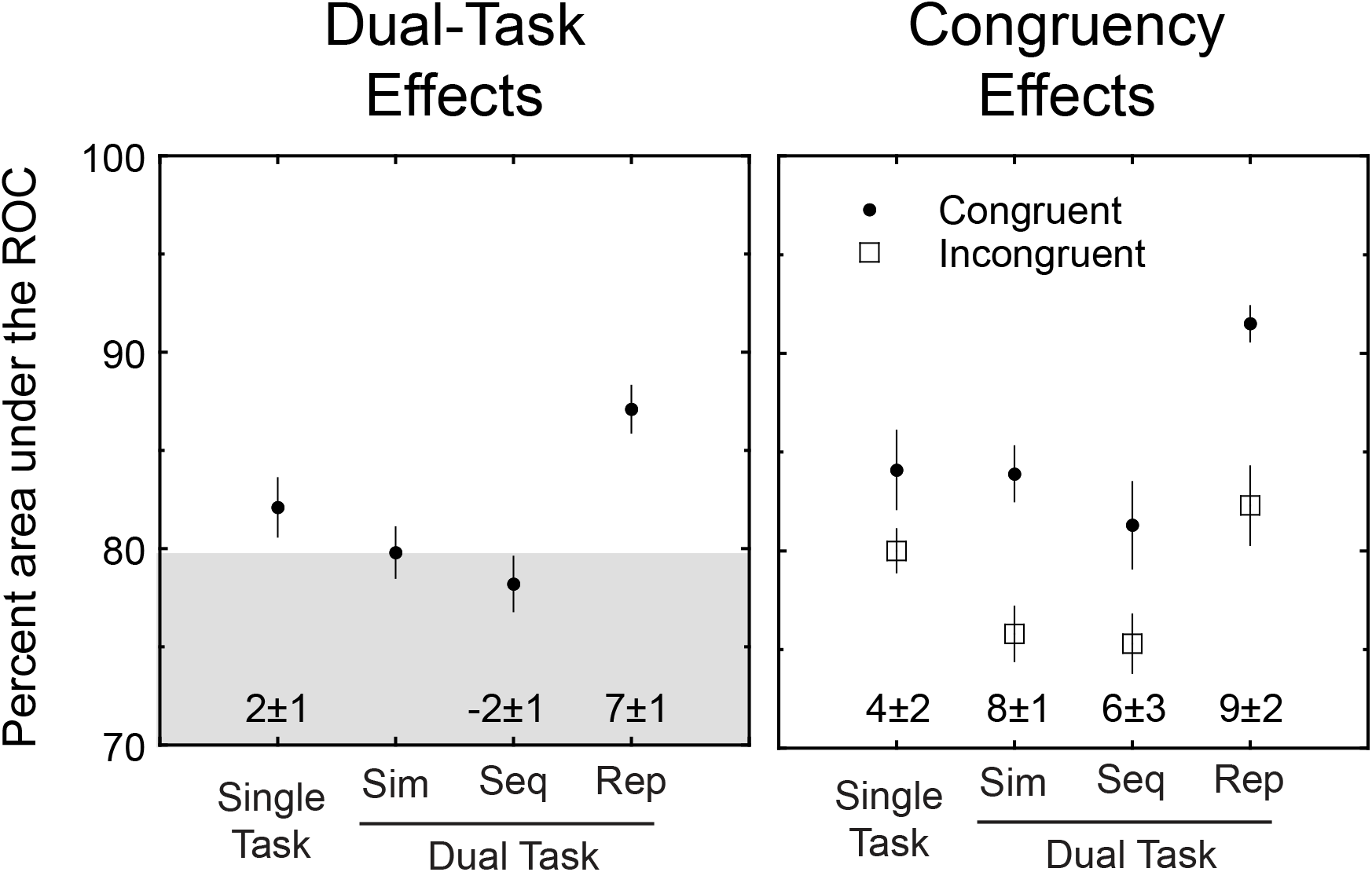
Results of Experiment 3. In the left panel, percent area under the ROC is shown for four conditions: single task, simultaneous dual task (sim), sequential dual task (seq), and repeated dual task (rep). The shading is anchored to performance in the simultaneous-dual-task condition to provide a standard of comparison. In this experiment, the single task and sequential dual task yielded similar performance to the simultaneous dual task. In contrast, the repeated dual task yielded better performance. This pattern of results is consistent with no dependency between the two tasks. In the right panel, percent area under the ROC is plotted for four conditions: single task, simultaneous dual task (sim), sequential dual task (seq), and repeated dual task (rep). These conditions are further broken down by whether the trial had congruent or incongruent stimuli. There were congruency effects for all conditions.

The dual-task deficit was 2.3±0.5% and was reliable for this experiment. By comparison, in Experiment 1 it was 1.6 ± 0.9% and only marginally reliable. An additional experiment described shortly will also show it to be about 2%. Thus, the experiments in this article are consistent with a dual-task deficit of about 2% for Gabor detection. This is small relative to the 5% deficit found for words, the 7% repeated effect, and the 8% dual-task deficit predicted by the fixed-capacity, parallel model (see Appendix B). To foreshadow our conclusions, the dual-task deficit is not completely absent for Gabor detection, but it is small relative to these other standards.

In the right panel of Figure 10, the four conditions are broken down by congruency and the values of each congruency effect are given at the bottom of the figure. There were reliable congruency effects in all conditions: the single-task congruency effect was 3.8±1.5%, (t(5)= 2.56, p<.05); the simultaneous congruency effect was 8.5±1.4%, (t(5)= 6.17, p<.001); the sequential congruency effect was 6.1±2.7%, (t(5)= 2.27, p<.05); and the repeated congruency effect was 9.0±1.8%, (t(5)= 5.14, p<.01). The single-task effect was smaller than the other effects. For example, it was half of the effect for simultaneous (3.8 vs. 8.5) and this difference was reliable (4.7±1.4, t(5)= 3.27, p<.05). Recall the congruency effects in Experiment 1 with similar stimuli were 8% for the dual-task condition and 2% for the single-task condition. Combining over both experiments there were 4 dual-task conditions that on average had a congruency effect of about 8% and two single-task conditions that on average had a congruency effect of about 3%. Thus, for Gabor detection there appears to be larger congruency effects for dual tasks relative to single tasks.

One aspect of these congruency effects deserves further scrutiny. For most of the conditions, the stimuli were presented simultaneously which is consistent with a congruency effect that depends on either immediate or later processing. The sequential task is different. Now the stimuli were presented sequentially with a full second elapsed between displays. If the congruency effect depends on immediate processing then it should be absent in the sequential condition. In fact, there was a reliable 6.1±2.7% congruency effect in the sequential condition. This result is consistent with the congruency effect being mediated by memory, decision or response processes rather than immediate processing. This result is an interesting contrast to the results found with words in the next experiment.

## Experiment 4

### Methods

In Experiment 4, we applied the simultaneous-sequential manipulation to word categorization. As with Experiment 3, there were 4 main conditions: single task, simultaneous dual task, sequential dual task and repeated dual task. After practice, six observers participated for 7 hours resulting in 672 trials in each of the 4 main conditions for each observer.

### Results

In the left panel of Figure 11, performance in terms of percent area under the ROC is shown for the 4 conditions. Performance was the worst for the simultaneous dual-task and better for the other three conditions. The shading emphasizes the simultaneous dual-task condition as a baseline upon which to compare the other conditions. The value of the difference from baseline are given at the bottom of the figure. The dual-task deficit (single-dual) was 5.2±0.7%, (t(5)= 7.54, p<.001); the sequential effect (sequential-simultaneous) was 5.3±0.8%, (t(5)= 6.51, p<.001); and, the repeated effect (repeated-simultaneous) was 9.9±1.1%, (t(5)= 9.15, p<.001). This is the pattern of effects expected for a dependency in immediate processing: the dual-task deficit and the sequential effect are similar. The fact that they are about half of the repeated effect shows that this dependency is less than expected from a fixed-capacity limit on processing.

**Figure 11.**
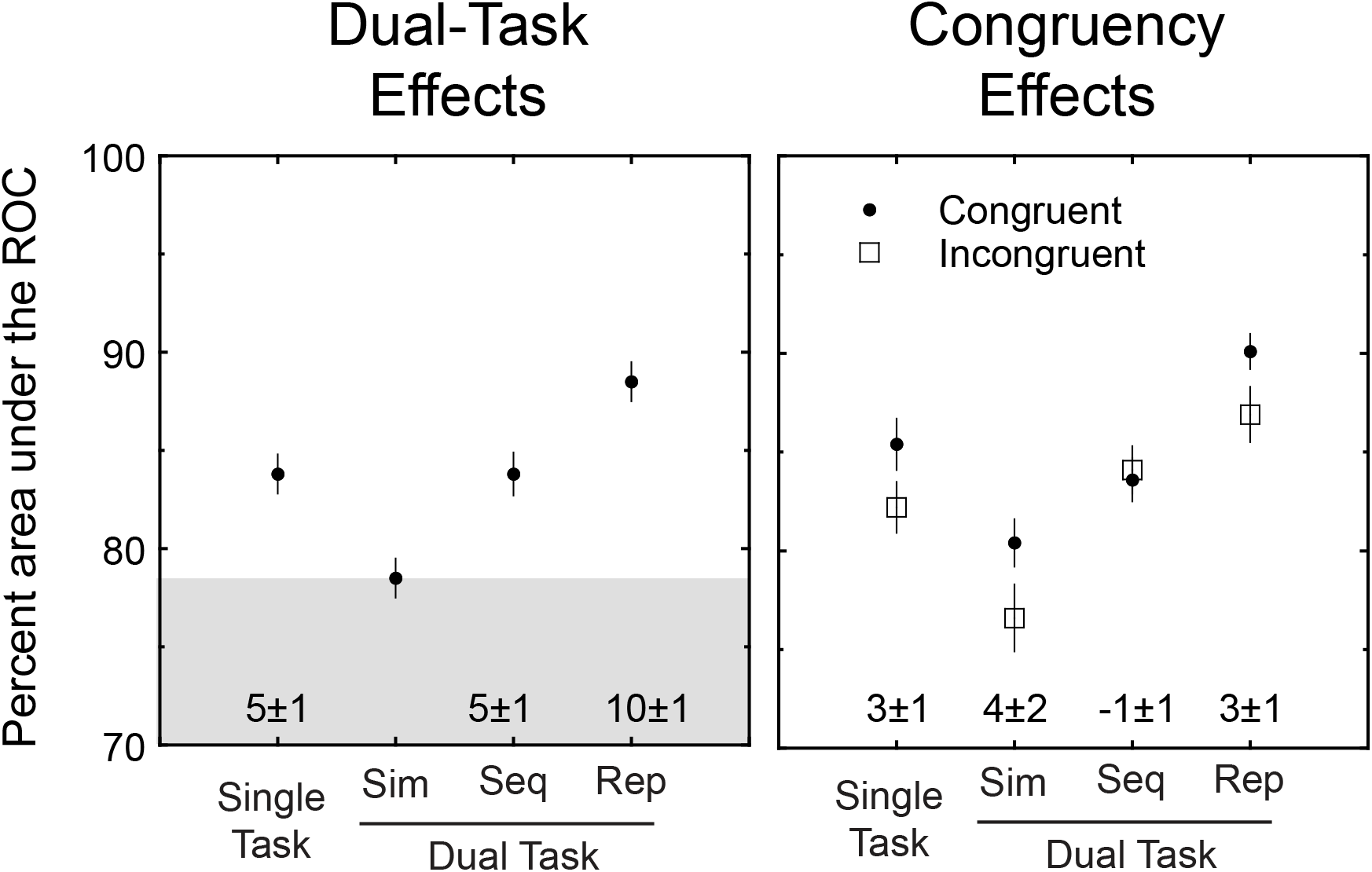
Results of Experiment 4. In the left panel, the percent area under the ROC is shown for four conditions: single task, simultaneous dual task (sim), sequential dual task (seq), and repeated dual task (rep). The shading is anchored to performance in the simultaneous-dual-task condition to provide a standard of comparison. In this experiment, the single task and sequential dual task yielded better performance than the simultaneous dual task. In addition, the repeated dual task yielded better performance than all of the other conditions. This pattern of results is consistent with a dependency in immediate processing between the two tasks. In the right panel, percent area under the ROC is plotted for four conditions: single task, simultaneous dual task (sim), sequential dual task (seq), and repeated dual task (rep). These conditions are further broken down by whether the trial had congruent or incongruent stimuli. There were congruency effects in several conditions but they were smaller than found in Experiment 3.

Again, the sequential-dual-task condition is worth further emphasis. In this experiment the dual-task deficit and the sequential effect had a similar magnitude. Separating the two stimuli in time eliminated the dependency. This is consistent with the dual-task deficit depending on immediate processing.

In the right panel of Figure 11, the four conditions are broken down by congruency and the values are given at the bottom of the figure. There were reliable (or marginally reliable) congruency effects for three of the conditions: the single-task congruency effect was 3.2±1.4%, (t(5)= 2.32, p<.05); the simultaneous congruency effect was 3.8±2.1%, (t(5)= 1.84, p<.1); and, the repeated congruency effect was 3.2±1.0%, (t(5)= 3.12, p<.05). In contrast, the sequential congruency effect was near zero and in the wrong direction at −0.5±0.6%, (t(5)= 0.71, p>.1). Recall the congruency effects in Experiment 2 with similar stimuli were about 1% for the both the single- and dual-task conditions. Combining over both experiments there was an average congruency effect of about 2%. Thus, for word categorization there was a smaller congruency effects than for Gabor detection. It also should be becoming clear there was a complementary pattern for the two measures of dependency: larger dual-task deficits for word categorization than Gabor detection but larger congruency effects for Gabor detection than word categorization.

## Experiment 5

In this experiment, we further pursued the Gabor detection task of Experiment 1. Specifically, we wanted to confirm that the congruency effects found with Gabor detection persist despite introducing factors intended to reduce dependencies between the responses.

### Methods

This experiment had two conditions: a single-task condition and a dual-task condition with just one response. The details were the same as Experiment 1 with the following modifications:

1. Separate keys were used for the two responses. Using a separate small keypad, the four keys on the left edge were assigned to the left-side task and the four keys of the right edge were assigned to the right-side task. For both tasks, the relevant four keys were arranged vertically and from bottom to top referred to the same confidence levels as in Experiment 1: likely-no, guess-no, guess-yes, likely-yes. This arrangement minimized Simon effects and eliminated decision errors in which one attempted to respond to one side when prompted to the other.
2. Observers were instructed to emphasize accuracy and take their time. To encourage that, the prompt following the stimulus display was delayed for 1 s instead of the 0.25 s in previous experiments.
3. The nature of independence between the two responses was discussed with each observer. Specifically, the two-by-two contingency table of possible stimuli for each task was explained and it was emphasized that they should make the two decisions independent of one another.

After practice, six observers participated for 7 hours resulting in 1344 trials in each of the 2 main conditions for each observer.

### Results

In this experiment, the dual-task deficit was 1.8 ± 0.8% which was reliable (t(5)=2.19, p<.05). This is a similar small effect as found in the prior two Gabor detection experiments. Our main interest is the effect of congruency which is shown in Figure 12 for the two main conditions with the values of each congruency effect given at the bottom of the figure. For the single-task condition, the congruency effect was 1.6 ± 1.5% which is not reliable (t(5)=1.04, p>.1). For the dual-task condition, the congruency effect was a reliable 4.3 ± 1.6% (t(5)=2.76, p<.05). Thus, the pattern of congruency effects was similar to the prior experiments using Gabor detection: a convincing congruency effect for the dual-task condition and a smaller effect for the single-task condition.

**Figure 12.**
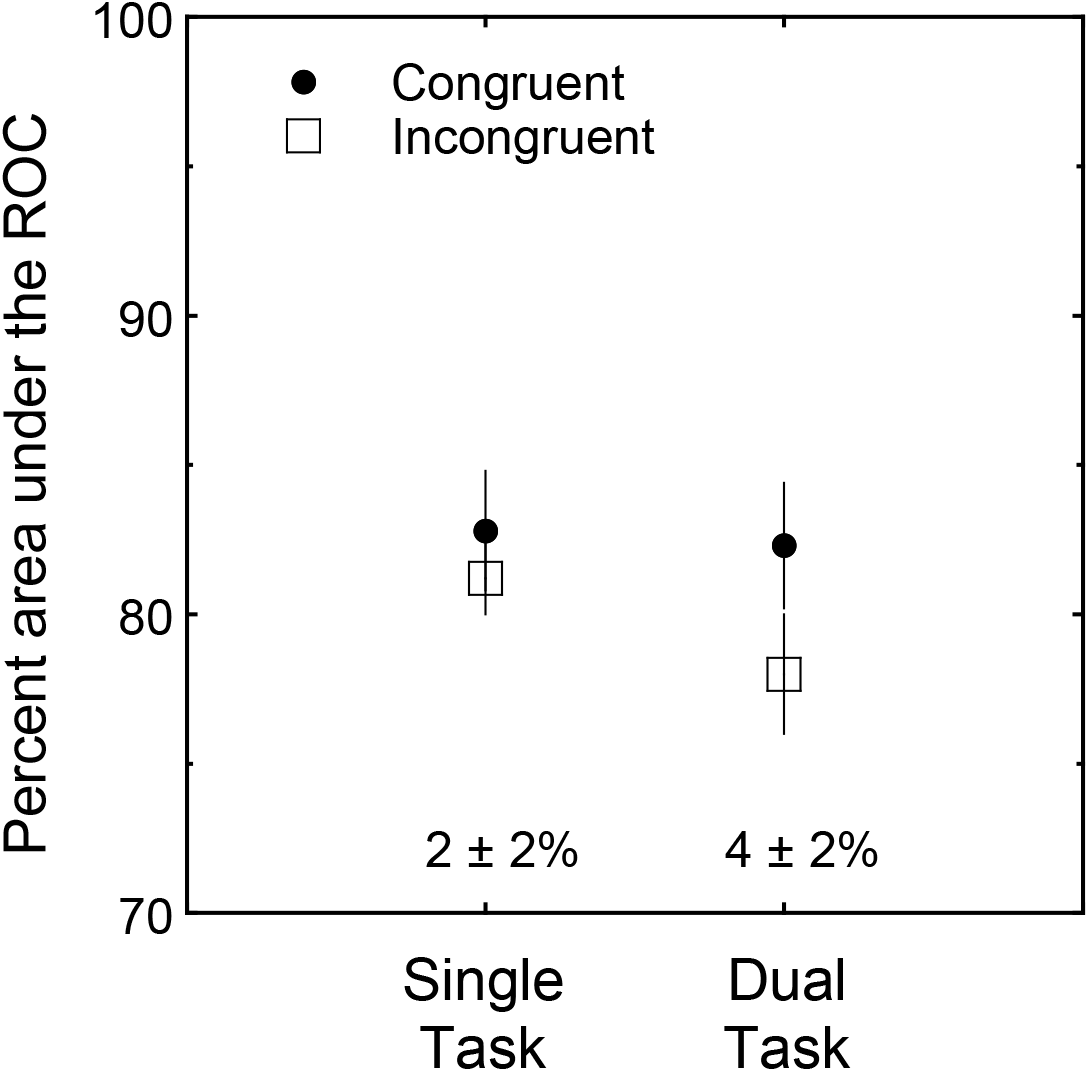
The congruency effects of Experiment 5. Percent area under the ROC is plotted for single-task and dual-task conditions. The congruency effects were reliable for only the dual-task condition.

## Experiment 6

In this experiment, we returned to word categorization. Unlike the Gabor detection experiments, word categorization judgments showed an effect of side. For example, in Experiment 1 in the single-task condition, performance on the left side was 83% and on the right side was 88% a difference of 5%. For the dual-task condition, performance on the left side was 75% and on the right side was 86% a difference of 11%. This advantage for the right side was found for every condition with words but not for Gabors. Here we asked if the observed dual-task effect is specific to conditions with unequal performance for the two sides. Specifically, we adjusted the contrast of the words on the left side to equate performance observed for the words on the right side. Will the dual-task deficit persist?

### Methods

In a preliminary experiment, the psychometric functions were measured for contrast using a single-task condition identical to that used in Experiment 1. For 5 observers, the psychometric functions appeared typical and thresholds for the left and right sides were 22 and 20% contrast, respectively. The ratio of the thresholds was reliably larger than one (1.09 ± 0.04, t(4) = 2.12, p<.05).

In the main experiment, we compared performance for single- and dual-task conditions (one response only) with an adjustment of contrast for the left and right sides. After practice, five observers participated for 5 hours resulting in 960 trials in each of the 2 main conditions for each observer. For three of the observers, the contrast was 22% and 20% contrast for the left and right sides respectively. For two other observers, the contrast was adjusted for their higher thresholds (24 and 22%; 32 and 27%). By accident, observer JH participated in only 1/3 of the intended number of trials. Will this roughly 2% difference in contrast eliminate the right-side advantage? And, if so, does the dual-task deficit persist?

### Results

In Figure 13, performance is shown for the single- and dual-task conditions broken down by left and right sides. With the contrast adjustment, there was no reliable difference between the sides (collapsed across conditions right - left = 1.0±1.6%, t(4)=0.62, p>.1). Combining across sides, the dual-task deficit was 5.2±1.4% (t(4)=3.78, p<.01). The congruency effects were also similar to before. For the single-task condition, the effect was 2.8±0.7% (t(5)=3.97, p<.01); and, for the dual-task condition, the effect was 3.2±1.4% (t(5)=2.24, p<.05). In sum, by adjusting contrast the right-side advantage was eliminated and the resulting dual-task deficit was still similar to the prior experiments.

**Figure 13.**
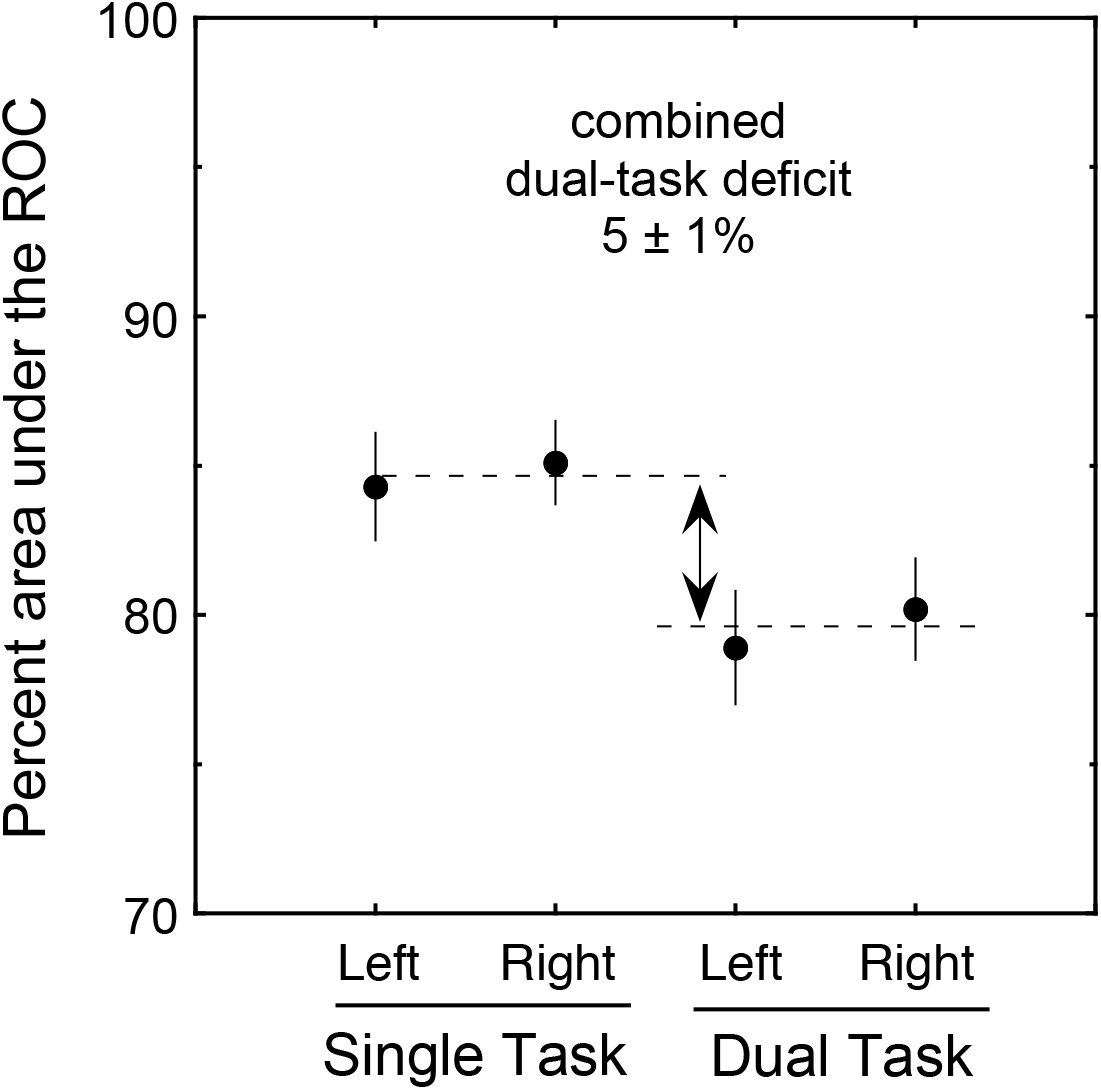
Results of Experiment 6. The percent area under the ROC is shown for single-task and dual-task conditions with a further breakdown by the left and right side. There was a reliable dual-task deficit regardless of side. In addition, for this experiment with a higher contrast on the left side compared to the right side, performance was similar for the two sides.

### Discussion

One cannot proceed without wondering why is there a sensitivity difference for words on the left side compared to the right side. This difference has been attributed to a several possibilities: hemispheric differences (e.g. Bryden, 1982), the reading order of English (Mishkin & Forgays, 1952), differential eccentricities of the critical first letters of the words (Kirsner & Schwartz, 1986), and scanning order preferences (Brysbaert, Vitu & Schroyens, 1996). The debate among these possibilities has not been settled.

## Experiment 7

For the standard serial model, dual-task deficits found with low contrasts are predicted to persist even with very high contrasts. This is because in the standard serial model, if a stimulus is not processed, the contrast is irrelevant and it will not be processed regardless of contrast. By comparison, for the parallel models, performance for high contrast conditions are predicted to reach near perfect levels and consequently eliminate the dual-task deficit. In addition, the same logic allows one to differentiate all-or-none models from graded models of congruency effects. We tested these high-contrast predictions for both Gabor detection and word categorization.

### Methods

This experiment included single-task and dual-task conditions for both Gabor detection and word categorization. The new feature was to use stimuli with 80% contrast rather than the 18-35% contrast used in the prior experiments. Otherwise, the details of the experiment follow those of Experiment 5 (e.g. separate keys for the left and right tasks). After practice, six observers participated for 4 hours resulting in 192 trials in each of the main conditions. All had previous experience in both Gabor detection and word categorization experiments.

### Results

For Gabor detection, the percent area under the ROC was 99.7±0.2% in the single-task condition and was 99.6±0.4% in the dual-task condition. The dual-task deficit was 0.2±0.2% and not reliable. In fact, four of the six observers were perfect on all trials of both conditions. Congruency effects were also very small and not reliable. Thus, for this judgment, performance was essentially perfect. As discussed below, this result rules any model of the dual-task deficit or congruency effect that predicts those effects persist for easily discriminable stimuli.

For word categorization, the percent area under the ROC was 97.8±0.9% in the single-task condition and was 95.9±0.8% in the dual-task condition. This yielded a small but reliable dual-task deficit of 1.9±0.7% (t(5)=2.55, p<.05). Congruency effects were small and not reliable. For this judgment, none of the observers were perfect. In the dual-task condition, observers made between 2% and 7% errors. In sum, the observed small dual-task deficit (2% instead of 5%) rules any model of dual-task deficits that has those effects persist unchanged or increased for easily discriminable stimuli.

## General Discussion

### Summary of Results

1. **Dual-task deficits**. For Gabor detection, there is a relatively small dual-task deficit while for word categorization there is a dual-task deficit that is intermediate between the magnitude predicted by parallel models of unlimited capacity and of fixed capacity.
2. **Dual-task congruency effects**. For Gabor detection, there is a dual-task congruency effect while for word categorization there is a smaller effect. These two results are illustrated in Figure 14. It is a scatterplot of the dual-task congruency effect plotted against the dual-task deficit. The three open squares illustrate the results of the three Gabor detection experiments and the three solid circles illustrate the results of the three word categorization experiments. Gabor detection has relatively large congruency effects (mean 6%) and relatively small dual-task deficits (mean 2%). In contrast, word categorization has relatively small congruency effects (mean 2%) and large dual-task deficits (mean 5%). These contrasting results were replicated in all three versions of each judgment.
3. **Single-task congruency effects**. For Gabor detection there is a smaller congruency effect in the single-task condition (mean of 2%) than in the dual-task condition (mean 6%). For word categorization, the congruency effects are similarly small for both single- and dual-task condition (means 2% and 2%).
4. **Sequential conditions**. For Gabor detection, the sequential condition of Experiment 3 has a similar dual-task congruency effect as found in the simultaneous condition. For word categorization, the sequential condition of Experiment 4 eliminates the dual-task deficit found in the simultaneous condition.
5. **High-contrast conditions**. For Gabor detection, the dual-task congruency effect observed for low contrast (mean 6%) essentially disappeared at high contrast (mean 0.5%). For word categorization, the dual-task deficit observed for low contrast (mean 5%) was reduced at high contrast (mean 2%).

**Figure 14.**
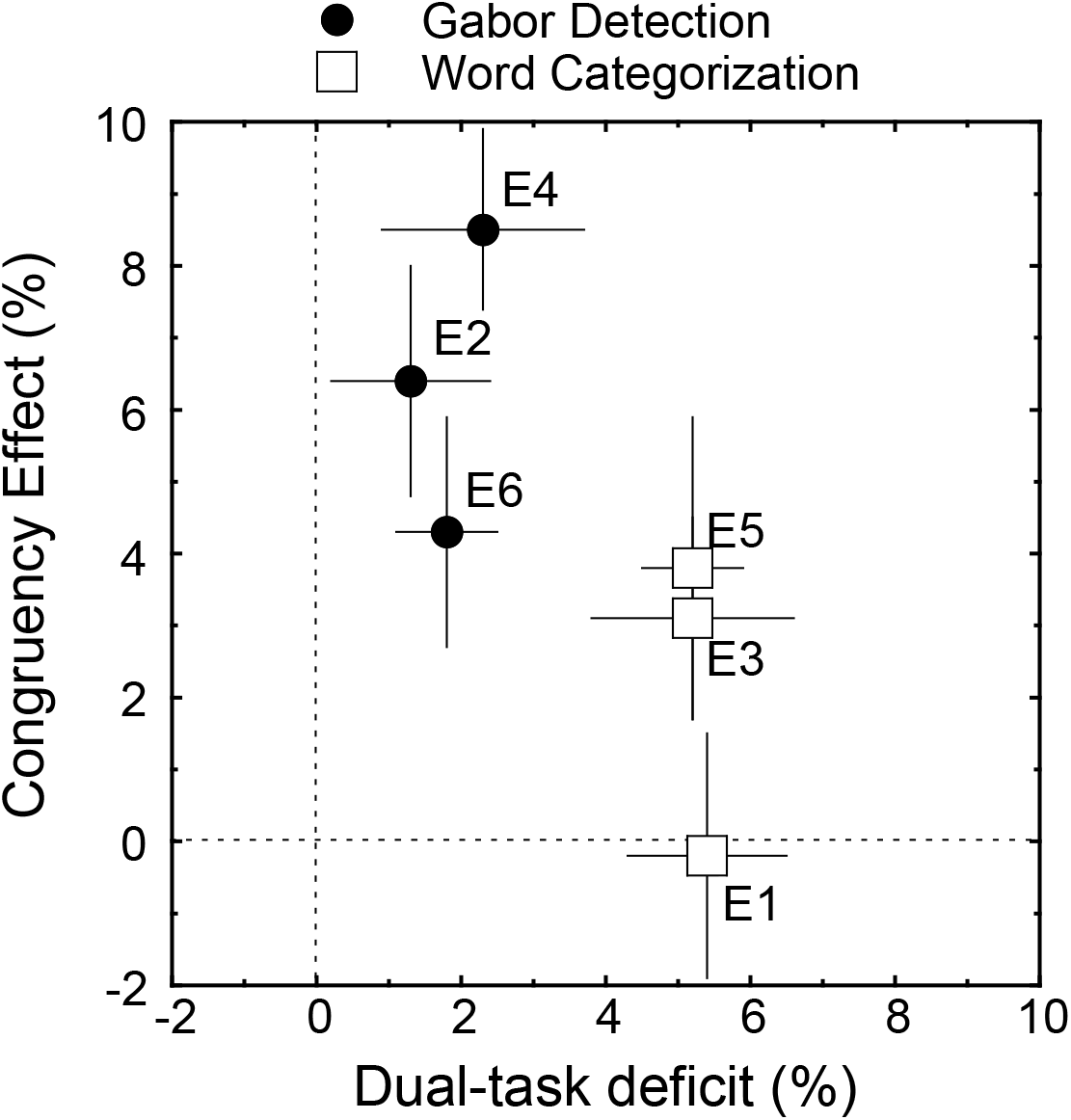
Summary of the main results. This is a scatter plot of the results of each experiment in terms of the dual-task deficit and the congruency effect found in the dual-task condition. The results for Gabor detection are shown by open symbols and the results for word categorization are shown by solid symbols. Word categorization had larger dual-task deficits than Gabor detection. In contrast, Gabor detection had larger congruency effects than word categorization.

### More than words versus Gabor patterns

It is easy to attribute the contrasting results to the use of words versus simple visual patterns. But that might be a mistake. These conditions also differed in other ways:

1. detection vs. categorization task,
2. specific targets vs. many possible targets, and
3. consistent mapping of stimuli and responses vs. variable mapping. We suspect that multiple factors contribute to the presence or absence of dual-task deficits and congruency effects.

### Working Hypotheses

#### Gabor detection

There are five results that constrain a theory for the Gabor detection task. The first result is the relatively small dual-task deficit. This is consistent with a limited-capacity, parallel model in which there is only a small limit on processing.

The second result is the congruency effect in the dual-task condition and not the single-task condition. This is consistent with selection error and not pure interactive processing.

The third result is that the congruency effect persisted with sequential presentation. This is consistent with the effect being due to later processes and not immediate processes.

The fourth result is that the congruency effect did not persist for high contrast stimuli. To analyze this result, we present two models in appendix B. One model has all-or-none selection (e.g. Palmer & Moore, 2009) and the other has graded selection (e.g. Kinchla et al., 1995). The high contrast results were consistent with the graded model and not the all-or-none model. To summarize the results up to this point, the results are consistent with graded selection in a later process such as decision.

The fifth result was unexpected. The response correlations were weakly positive which is generally consistent with parallel rather than serial models. The twist was that the positive correlation was specific to the noise-noise condition. To account for this result we modified the weighting model to give weight to noise trials and not signal trials. This modified model can yield the observed pattern of response correlations.

Stepping back from this specific model, one can also consider a hybrid theory that combines elements of selection error and interactive processing. In particular, suppose selection can occur early in the single-task condition but must occur late in the dual-task condition. Then early selection can be used to protect the stimulus representation from interactive processing that occurs in memory or response. Using this approach one can make the selection process be all-or-none and the interaction be graded.

In summary, we have just defined two accounts for the five results observed with Gabor detection. Moreover, we can reject many other hypotheses. These are summarized by the table of possible hypotheses in Figure 15. It elaborates the earlier 2 by 2 table by including hybrid hypotheses along with each pure hypothesis. Each hypothesis that can be rejected is marked by a red X and the two working hypotheses are circled in green. In addition, there are several other hybrid hypotheses that are also still viable. Of the original 2-by-2-by-2 table all but one of the original eight hypotheses can be rejected.

**Figure 15.**
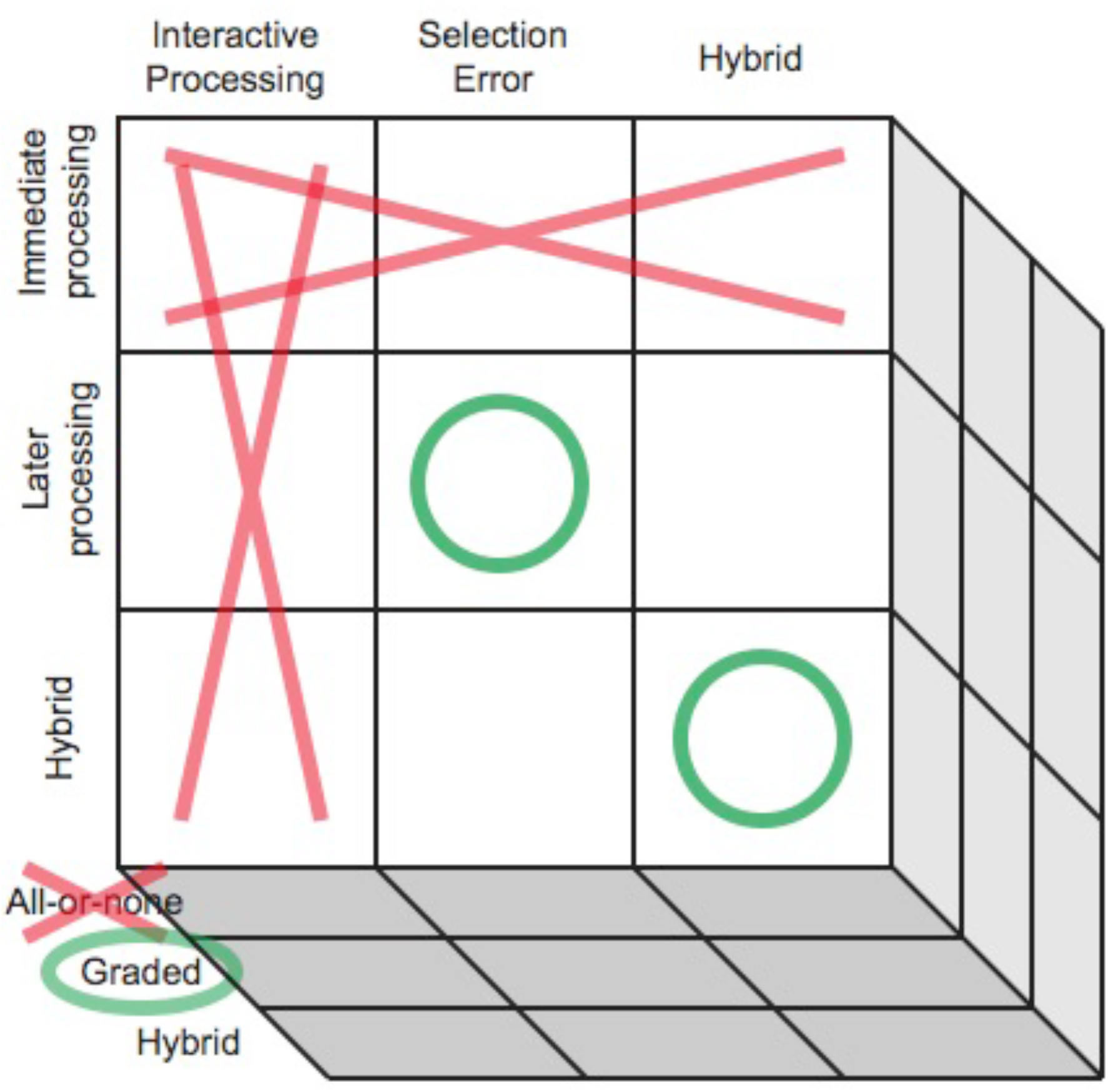
A summary of the theories considered in the analysis of congruency effects in Gabor detection. It is organized by the three distinctions introduced previously. The experiments rule out the theories marked by the red Xs. The two working hypotheses described in the text are marked by green circles.

#### Word categorization

Given the small size of the congruency effects for word categorization, there are fewer convincing results to distinguish among the alternative theories. The first result is a dual-task deficit that is consistent with a limited-capacity, parallel model. The second result was little sign of the negative response correlations expected from serial models. The third result was the reduced but not eliminated dual task deficit for the high contrast condition. All three of these results are consistent with the limited-capacity, parallel model. But it is also possible to account for these results by taking some modest liberties with the standard serial model. Thus, while there is no significant evidence for serial models, one cannot rule them out.

We urge caution in interpreting the generality of these working hypotheses. Our reading of the literature is that the variety of congruency effects will require a variety of models. For example, spatial filtering experiments with small separations between stimuli require a different theory than dual tasks with wide separations (Palmer & Moore, 2009).

### Relation to the Prior Literature

#### Gabor detection and dual-task deficits

Finding little or no dual-task deficit has several precedents for detection or detection-like judgments (e.g. Bonnel, et al., 1992). It is also consistent with parts of the visual search literature. In particular, the simultaneous-sequential paradigm has been used in visual search to separate effects of immediate processes such as perception from effects of later processing such as decision. For judgments of simple features, several studies have demonstrated no effect of the number of simultaneously displayed stimuli when the number of decisions was held constant (e.g. Huang & Pashler, 2005; Scharff et al., 2011). For example, Scharff and colleagues studied visual search for contrast increments and showed that simultaneously displays of 4 stimuli or two sequential displays of 2 stimuli resulted in similar performance.

These results conflict with claims that all visual task have limited capacity (e.g. Kahneman, 1973; Posner, 1980). One resolution to this conflict is that some prior tests of these claims did not specifically examine the case of detection or detection-like tasks. Another possible resolution is that some of the details of the procedure are important to obtaining a result of no deficit.

A more subtle argument is that all tasks have limited capacity to at least some small degree. Indeed, examining all three Gabor detection experiments reveals a mean dual-task deficit of about 2%. Thus, it seems likely that there is a small deficit for the procedure used here. Our response to this argument is to emphasize that there are clear differences in the magnitude of the dual-task deficit for different tasks that range from near zero to that predicted by a fixed-capacity model. This argument uses the prediction of fixed capacity as a reference point for what is a significant magnitude of the dual-task deficit. There is little doubt that the Gabor detection task has dual-task deficits closer to zero than to that predicted by fixed capacity (about 8% deficit for the current conditions). This contrasts word categorization which have dual-task deficits closer to the fixed-capacity prediction than zero. Thus, some judgments yield deficits that are closer to fixed capacity than unlimited capacity and other judgments yield deficits are closer to unlimited capacity.

#### Gabor detection and congruency effects

We also find dual-task congruency effects with Gabor detection. Such congruency effects were previously found by Bonnel et al. (1992) and Ernst et al. (2012). Considering more cognitive judgments and speeded dual-task paradigms, one finds that such dual-task congruency effects are common but perhaps not universal (e.g. Hommel, 1998; Logan & Gordon, 2001). Our result of finding a congruency effect with sequential presentation is consistent with mediation by later processes such as memory, decision or response. It is not consistent with mediation by immediate processes such as perception or memory encoding. Thus, it seems likely that this kind of dependency does occur but, at least for widely separated stimuli, has more to do with later processing than immediate processing.

Unlike the dual-task conditions, there was little congruency effect for the single-task conditions. A similar difference has been reported for an auditory detection task (McCloy & Lee, 2015). This divergence has been studied in the more cognitive speeded dual-task literature (e.g. Logan & Gordon, 2001). More generally, measuring congruency effects in the single-task condition is analogous to measuring congruency effects in the flanker paradigm with wide separations between the targets and flankers. In the flanker paradigm it is widely reported that increasing the separation decreases the congruency effect (Eriksen & Eriksen, 1974; Yantis & Johnston, 1990). Thus, it appears that a large separation and foreknowledge of the relevant stimuli are sufficient to prevent congruency effects with an irrelevant stimulus. This conflicts with claims that interactive processing is widespread at all levels of processing (Navon & Miller, 1987). Instead, it is consistent with either selection errors or interactive processing that can be avoided in the single-task condition.

More theoretically, numerous theories include the idea that perceptual processing of simple features occurs independently under ideal circumstances such as widely separated stimuli. In the vision literature, there are theories of spatial vision that tile the visual field with independent and parallel processing channels (e.g. Graham, 1989). In the visual search literature, there are several “two-stage search theories” that have a first stage with independent processing of well separated stimuli (e.g. Hoffman, 1978; Treisman & Gelade, 1980). In the selective attention literature, there are theories that presume interactions for nearby stimuli that disappear for widely separated stimuli (Palmer & Moore, 2009; Yantis & Johnston, 1990). The current results with no dual-task deficit and congruency effects attributed to later processing are consistent with such independent perceptual processing of widely separated stimuli.

#### Word categorization and dual-task deficits

Finding a dual-task deficit for a pair of word categorization judgments has several precedents in the literature. For auditory word identification tasks it has been extensively reported (e.g. McCloy & Lee, 2015). Surprisingly, it has only recently been measured with an accuracy dual task in vision (White et al., 2018 described below). Previously, it had been measured with a “secondary task” approach to dual tasks (e.g. Becker, 1976; Herdman, 1992) and by the “slack logic” approach to perceptual dependencies using speeded dual tasks (e.g. McCann, Remmington & van Selst, 2000). The current results using accuracy dual tasks complement these other approaches and provide relatively direct support for a perceptual interpretation of the dependency. This interpretation is further supported by the finding that the dual-task deficit for words is eliminated by a sequential presentation. In sum, the present study joins the prior literature in suggesting limited-capacity processing for the immediate perception of two words.

This interpretation is further supported by studies of words in visual search. While set-size effects in visual search have multiple interpretations (Scharff, et al., 2011; Townsend & Ashby, 1983), the simultaneous-sequential comparison in visual search tests specifically for a dependency in immediate processes such as perception or memory encoding. Several studies have shown that judging multiple words is better for a sequential presentation compared to simultaneous presentation (e.g. Harris, Pashler & Coburn, 2004; Scharff, et al., 2011). A particularly nice demonstration is in the dissertation of Patterson (2006). She replicated a study by Pashler and Badgio (1987) that had previously shown that judging the highest digit did not show a sequential advantage for character displays. Patterson replicated this result and then tried it with words. When words were used instead of characters (“seven” instead of “7”), there was a sequential advantage. Thus the effect appears to be specific to words and not single characters that convey the same information.

These results are not consistent with two words being processed in parallel with unlimited capacity. Such a result is predicted by the strong automaticity hypotheses that predict both inflexible processing and unlimited capacity for automatic processes. Such a view is most common as an account of the Stroop phenomena (e.g. Brown, Gore & Carr, 2002) as a failure of selection due to inflexible processing of all words. Such a view has been challenged by studies that directly manipulate the divided attention aspect of the theory by varying the number of words in a Stroop task (Kahneman & Chajczyk, 1983). Increasing the number of words decreases the size of the Stroop effect which has been called dilution and attributed to limited capacity processing of the words. Related arguments have also been marshaled in other variations on the Stroop paradigm (Stolz & Besner, 1999; Risko, Stolz & Besner, 2005). Thus, while the Stroop paradigm is clearly an example of the failure of selection, it is probably not an example of the unlimited-capacity processing of words. In sum, the dilution effect and related phenomena have undermined the most common argument for words being processed with unlimited capacity. The experiments presented here reinforce the case against words being processed with unlimited capacity.

The results reported here are also not what is predicted by theories with strictly serial processing of two words in word recognition. They also differ from what is predicted by theories of serial processing in reading (Reichle, Pollatsek, Fisher & Rayner, 1998, but see Engbert, Nuthmann, Richter & Kliegl, 2005). As discussing previously, a simple all-or-none serial model predicts large-magnitude dual-task deficits and a negative correlation in the trial-by-trial accuracy of the responses to the two words. These effects were not found in the current studies. For an alternative and perhaps more general test of serial models, one can use the redundant target paradigm (e.g. van der Heijden, 1975). Indeed, Mullin and Egeth (1989) found support for the serial processing of words but the study of Shepherdson and Miller (2014; 2016) came to a contradictory conclusion.

Evidence that something is amiss in our story comes from a recent study from our lab that replaced the simultaneous noise used in the current study with post masks (White, et al., 2018). In that study, we found the larger dual-task deficits and the negative correlations predicted by the all-or-none serial model. That study builds on previous work suggesting that post masks yield different results in attention paradigms than found with simultaneous noise or paradigms without any kind of noise or mask (Morgan, Ward & Castet, 1998; Smith, Ellis, Sewell & Wolfgang, 2010). Why the difference? One possibility is that masking terminated processing and prevented switching from one word to the other before the memory of the second is lost. That is consistent with a serial model of word processing. Another possibility is that masking changed the processing in more general ways (e.g. Eriksen 1980). For example, the post masks might have caused object substitutions that are themselves dependent on where selection is directed (Enns & Di Lollo, 1997; Enns, 2004). That is not consistent with a serial model of word processing outside of masking. In sum, the evidence for serial or parallel processing in words depends on the details of the experiment.

#### Word categorization and congruency effects

For word categorization, there were relatively small congruency effects. This is for both single-task conditions and dual-task conditions, and is in contrast with Gabor detection where there were larger congruency effects for the dual-task conditions. To our knowledge, the most similar result are the reduced congruency effects found in audition by McCloy and Lee (2015) for semantic versus phonetic tasks.

In vision, perhaps the closest precedents are flanker studies that use pairs of words. Several studies have presented pairs of words near each other and cued one as relevant and the other as irrelevant. Under these conditions, they typically observe congruency effects (Dallas & Merikle, 1976; Shafer & LaBerge, 1979; Lambert, Beard & Thompson, 1988). This is unlike the current study with widely separated words. Furthermore, a few studies have shown that increasing the distance between the words decreases congruency effects (e.g. Broadbent & Gathercole, 1990). Thus, it seems likely that one of the reasons our study did not show congruency effects is because the words were widely separated.

Another more distant precedent are masked priming studies that present two words in sequence with one word serving as a potential prime of the other target word. Once again, congruency effects are observed for words close together in time and space. For example, Lachter et al. (2004) manipulated the spatial and temporal relation between the prime and target word. Congruency effects were reduced for more widely separated words similar to the result in the flanker literature. Based on several additional manipulations, they argued that congruency effects occur under conditions in which the prime is attended rather than ignored as observers were instructed. Again, our experiments provide good conditions for selecting the relevant word and ignoring the irrelevant word with the result of little or no congruency effect.

Both of these precedents speak to the single-task condition in our experiments. They say less about why there is no congruency effect in the dual-task conditions in which both words are relevant to the task. The lack of congruency effects for widely separated words is challenging for theories that assume widespread interactive processing of words. We take the view that such interactive processing does occur but is specific to particular domains such as crowding and response selection.

### Implications for General Theories of Divided Attention

In the introduction, we outlined four general theories of divided attention: theories with a single resource, theories with interactive processing, theories of automaticity, and theories with multiple dependencies. What can be said about them based on the results of this study?

One of our results is that dual-task deficits and congruency effects do not necessarily covary. This is inconsistent with any single dependency theory. In addition, only the Gabor detection task showed robust congruency effects. Thus these effects are not universal. Furthermore, the congruency effect observed for Gabor detection is probably not mediated by immediate processing because it persists with sequential presentation. Thus, there is little or no evidence of immediate interactive processing in the perception of widely separated stimuli. This result is admitted particular to our situation. If one considered closely separated stimuli there would have been interactions (Parkes et al., 2001); Or, if one introduced contingencies, there would have been interactions (Mordkoff & Yantis, 1991). Those are situations known to yield interactions. We argue that while interactive processing occurs in some situations, they are not universal and cannot underlie the entire variety of divided attention effects.

Our results are consistent with theories of automaticity: Gabor detection depends on an automatic process that avoids dual-task deficits at the cost of congruency effects. In contrast, word categorization depends on a controlled process that avoids congruency effects at the cost of dual-task deficits. This is a possible account. But, we suspect on the basis on ongoing research that there are also situations with both effects, and other conditions with neither effect. If so, such results are harder to account for by theories of automaticity.

Instead, we argue for the more elaborate theories that include multiple modules which have different dependencies. With respect to perception, there was limited capacity for word categorization but not Gabor detection. There were no congruency effects for our widely separated words, but it is likely that such congruency effects would be common with more closely spaced stimuli (crowding). Thus, for perception, there are capacity limits for some situations (words and objects) but not others (widely separated simple features) and there is interactive processing for closely spaced but not widely spaced stimuli. The variety of dependencies depends on the task and on space. We suspect a similar variety of dependencies exists for other modules such as memory and response processing.

This view is similar to the view presented in Pashler (1989) who made the case for separate dependencies in perception (stimulus identification) and in response (response selection). Since that work, others have asked if other perceptual phenomena follow the same pattern. For the attentional blink, Ruthruff and Pashler (2001) found evidence it is related to the response dependency rather than the perceptual dependency. For multiple object tracking, several studies have found evidence it is also related to the response dependency (Tombu & Seiffert, 2008) and not the perceptual dependency (Kunar, Carter, Cohen & Horowitz, 2008). Thus, there is growing evidence consistent with multiple dependencies causing effects of divided attention. We add to this account the existence of multiple loci of interactive processing and failures of selection which contribute to effects of both selective and divided attention.

## Conclusion

There were different patterns of divided attention effects for two judgments. For Gabor detection, there were dual-task congruency effects but little dual-task deficit. The simultaneous-sequential experiments showed that the congruency effects were not specific to immediate processing and probably instead due to later processes such as memory, decision or response. Further results were consistent with theories for the congruency effects that dependent on either errors in a graded selection process or an all-or-none selection process that can protect against graded interactive processes in the single-task condition but not the dual-task condition. These results are consistent with independent processing in perception for widely separated simple stimuli. For word categorization, there were dual-task deficits but little congruency effect. The simultaneous-sequential experiments showed that the dual-task deficit was specific to immediate and not later processing. These effects can be modeled by limited-capacity, parallel processes in perception but we cannot rule out serial processing. Together, this set of results rule out single dependency theories of divided attention. Instead, the results are consistent with different dependencies in perception and later processes.

## Appendix A

## Direct Analysis of the Rating Data

In the body of this article, we summarized performance by *A*_*ROC*_: the area under the ROC curve. As discussed in methods, this measure is a rough and ready estimate of the unbiased proportion correct. In this appendix, we describe the rating data from which *A*_*ROC*_ is estimated and consider the issue of response bias.

## Dual-task effects

Rating data such as used here can be summarized by an ROC graph that plots the percent hits against the percent false alarms. Figure 16 shows such parametric plots for each of the six main experiments. In each panel, there are separate symbols for the single-task (solid circles) and the dual-task condition (open squares). Such graphs represent the cumulative percent of responding “yes” for each consecutive rating (rating 1, rating 2 or less, rating 3 or less). There is no point for the fourth rating because one must use one of the 4 ratings. If performance is at chance, the points will fall on the positive diagonal; if performance is perfect, it will fall in the upper left corner with 100% hits and 0% false alarms. Finally, if performance for a given rating is unbiased (probability of a “yes” is 0.5), then that point will fall on the negative diagonal.

**Figure 16.**
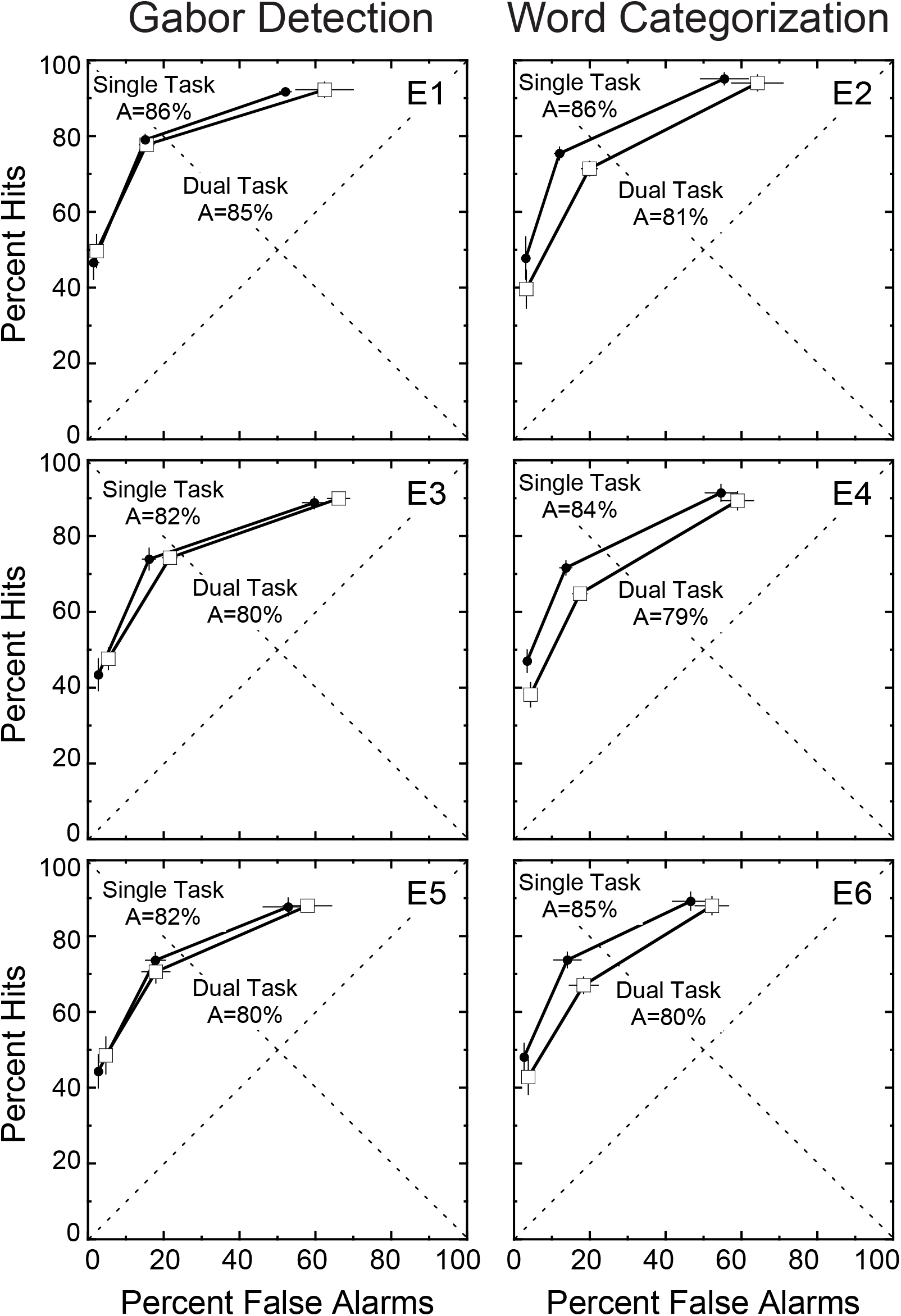
ROC functions for single-task versus dual-task conditions. The results of the six main experiments are shown in separate panels. In each panel is an ROC plot in which the percent hits is plotted against the percent false alarms. Single-task conditions are shown by the solid symbols and dual-task conditions are shown by the open symbols. There were effects of the dual task for all of the word categorization experiments and smaller effects for the Gabor detection experiments. This corresponds well with the analysis in the body of the article using estimates of the area under the ROC function.

For all experiments, the three points on the ROC curve formed a negatively accelerated function that is typical of predictions from signal detection theory based upon comparing random variables to a decision criterion (Green & Swets, 1966). It clearly deviated from a linear function that is predicted by the high threshold theory (a line from (0,x) to (100,100) with x between 0 and 100) in which one guesses when not detecting the target. Thus, one can rule out a simple version of the high threshold model for both experiments.

For Gabor detection, the points for single- and dual-task conditions tended to fall on top of one another which is also reflected in percent *A*_*ROC*_ having nearly identical values. Little or no dual-task deficit.

For word categorization, the points that form the two ROC functions were separated. This was captured in the reliably different percent *A*_*ROC*_ values for the two conditions. Thus, the dual-task deficit was clear in the rating data itself as well as the summary *A*_*ROC*_ measure.

Now consider whether response bias is different for the single- and dual-task conditions. A common way to summarize response bias is to use the middle point of the ratings which was instructed to differentiate between a “no” and a “yes” response. In other words, this point is an estimate of the proportion of “yes” responses given a target (hits) and given a distractor (false alarms). These two values can be combined to give an overall proportion of “yes” responses which provides a measure of response bias. In all of the conditions, this measure was near 45% which means the observers were conservative about making a “yes” response (falls below the unbiased line). For example in Experiment 1 (Gabor detection), the percent “yes” responses was 47.0 ± 0.8% for the single-task condition and was 46.6 ± 1.4% for the dual-task condition: an unreliable difference of 0.4 ± 1.6%. Over all experiments, the percent “yes” was 44.8% for the single tasks and was 44.9% for the dual tasks. Thus, there was no sign of an effect of dual tasks on response bias.

## Congruency effects

The effect of congruency on the ROC is illustrated in Figure 17 using similar conventions as the previous figure. Each panel shows for one experiment the results of the dual-task condition broken down by congruent and incongruent trials. Gabor detection results are shown on the left and word categorization results are shown on the right. Only the dual-task conditions are shown because the congruency effect is larger for that condition than the single-task condition.

**Figure 17.**
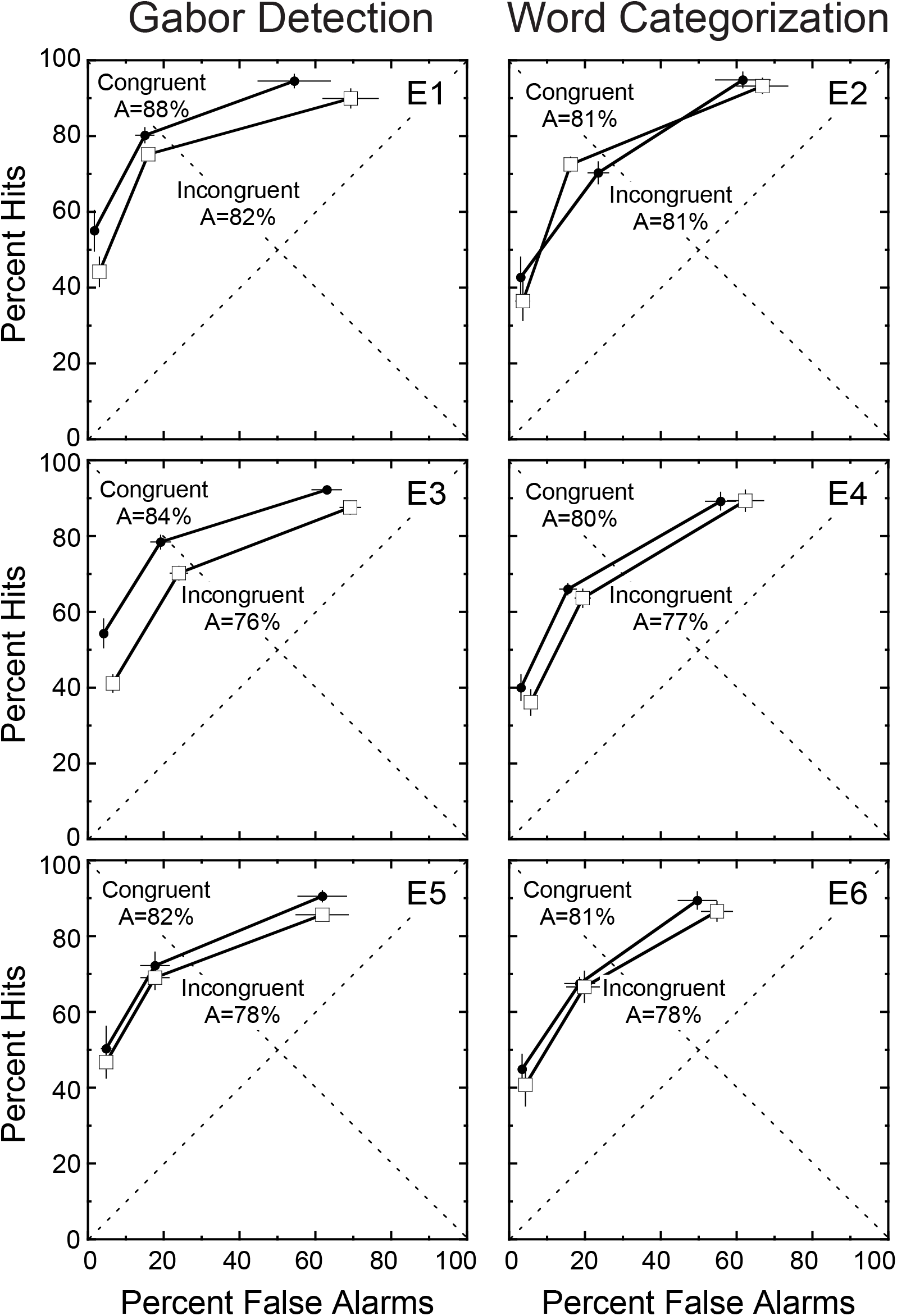
ROC functions for congruent versus incongruent conditions. The results of the six main experiments are shown in separate panels. In each panel is an ROC plot in which the percent hits is plotted against the percent false alarms. Congruent conditions are shown by the solid symbols and incongruent conditions are shown by the open symbols. There were effects of congruency for all of the Gabor detection experiments and smaller effects for the word categorization experiments. This corresponds well with the analysis in the body of the article using estimates of the area under the ROC function.

For the Gabor detection experiments, the congruency effects were consistently reliable as reported in the body of the article. Here one can also see the ROCs were as one would expect for a shift of sensitivity between the congruent and incongruent conditions. For the word categorization experiments, the congruency effects was smaller and not always reliable for individual experiments.

The effect of congruency on response bias can be summarized by the congruency effect on the percent “yes” responses as before. The congruency effect on percent “yes” varied from - 0.8% to 2.5% across experiments with a grand mean difference of 1.2%. This difference was reliable for only 1 of the 6 experiments. Thus there was little evidence for an effect of congruency on response bias.

## Averaged ROC functions

The graphs shown in this analysis are the mean of the observers in each experiment. Thus, one can ask how representative are these mean graphs of the graphs for individual observers. The short answer is quite representative. The individual graphs all share the same shape of a negatively accelerated function. They also have central points in similar locations and thus have similar values for the percent “yes” responses. This can be inferred from the small error bars for the central point. Where individuals do vary is how extreme are the other two points. Different observers vary considerably in how often they use the more extreme ratings. This can be seen by the relatively large error bars for one of the dimensions of the extreme points. For example, for the more “liberal” rating in the upper right corner, the horizontal error bars are quite large. Individuals vary in the location of these points and the error bars for individual observers are much smaller than the error bars of the mean data. In summary, the mean data is representative of the shape of the function and the overall response bias. But individuals do differ in how they use the endpoints of the rating scale.

In sum, the *A*_*ROC*_ measure succeeds in capturing the differences and lack-of-differences seen for the dual-task and congruency conditions as shown by the ROC functions. The ROCs do not change in shape and there are no systematic effects of the manipulations on response bias.

## Appendix B

## Definition of the Limited-Capacity, Parallel Model

For our initial model, we assume a limited-capacity, parallel process. This will be developed via its special cases of unlimited-capacity and fixed-capacity models. Denote the two tasks: *i* = 1, 2. Assume the relevant evidence from each stimulus corresponds to random variables *S*_*1*_ and *S*_*2*_. Further assume these of these random variables are independent and identically distributed. For the tasks considered here, stimuli are either targets or distractors. For Gabor detection the targets were Gabor patterns in noise and the distractors were noise alone. For word categorization, the targets were words from the relevant category and distractors were words from any other category. Assume the difference between target and distractor representations corresponds to a shift in the mean of their random variables: *S*_*target*_ = *S*_*distractor*_ + *d*. The *sensitivity* parameter *d* is in units of the standard deviation of these random variables.

## Capacity limits

To begin, consider a fixed-capacity, parallel model as implemented by Shaw’s (1980) sample size model. In Shaw’s model, the variability of the representation is inversely proportional to the number of samples assigned to it. Divided attention is allocated by distributing a fixed number of samples among the relevant stimulus representation. Let the initial sensory representation be denoted by the random variable *S*\*. Then if *m* samples a drawn from *S*\* the resulting representation *S’(1)* is the mean of the samples:

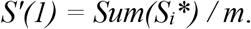

Assume one allocates some fraction *a* of the *m* samples to one stimulus and that 0 ≤ *a* ≤ 1. The resulting representation *S’(a)* is

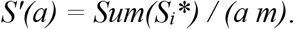

For example, if there were *n* stimuli and one equally sampled the stimuli the result is *S’(1/n)*. The consequence of combining independent samples additively is that the variance of S’(a) declines proportionally with the fraction sampled:

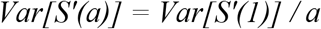

(for the derivation see Palmer, Ames & Lindsey, 1993).

Here we normalize the variance and scale discriminability accordingly. Now the relevant term is the standard deviation that varies with the square root of *1*/*a*. Thus, if the random variable is scaled by Sqrt(1/*a*) to normalize it, then the discriminability parameter must also be scaled by the same factor. Lets denote the discriminability parameter for a single stimulus as *d(1)*. Then one can represent the discriminability with *a* samples as *d(a)* where

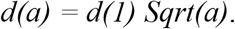

In sum, the attentional parameter *a* expresses the amount of processing (proportion of samples) allocated to each stimulus. A value of 1 indicates full processing as assumed for a single stimulus; a value of 0.5 indicates an equal split of processing as assumed for two relevant stimuli; and a value of 0 indicates the stimulus is ignored.

## Decision rule

Assume the decision rule for task *i* is:

If *S′*_*i*_ > *c* then respond “yes”, otherwise, “no”.

Following this rule, the proportion of “yes” responses for task *i* is:

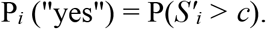

The *criterion* parameter *c* allows one to adjust the response bias for the decision. It is also in units of the standard deviation of the random variables. In fact, for the rating task one actually needs three criteria to generate 4 possible ratings, but we ignore that modest complication in this appendix.

## Generalization to “limited capacity”

To generalize this model we add an additional parameter, *k*, to express the degree of limited capacity. We want *k* = 1 to yield the just described fixed-capacity model. In addition, we want *k* = 0 to yield an unlimited-capacity model and intermediate values of *k* to represent varying degrees of limited capacity. This can be done building on the sample size model by defining the effective number of samples as a Minkowski metric of the individual samples: the effective proportion of samples is defined by *a*^*k*^. Using this definition, the effect of attention allocation becomes

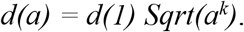

For *k* = 1, the effect of a remains unchanged and the model has fixed capacity. For *k* = 0, then *a*^*k*^ = 1 for any *a* and the model has unlimited capacity. For intermediate values of *k*, the number of samples is inflated relative to the fixed-capacity, parallel model. For example, for *k* = 0.5. An allocation of attention (*a* = 1/4) such as one might do over four stimuli results in *a*^*k*^ = 1/2. The effective number of samples is reduced to a smaller degree than occurs with *k* = 1. Finally, one can also use values of *k* > 1 to describe dependencies larger than expected from fixed capacity.

In summary, this limited-capacity, parallel model has four parameters: *d* is the sensitivity parameter, *c* is the decision criterion, *a* is the attentional parameter and *k* is the parameter for the degree of limited capacity. As described below, some of these parameters can be eliminated for some predictions.

## Alternative Models for Congruency Effects

For congruency effects, the coarsest contrast is between hypotheses with independent processing and perfect selection that predict no congruency effect and those with either selection errors or interactive processing that do predict a congruency effect. As described in the body of the article, the relevant hypotheses can be characterized on three dimensions: selection errors versus interactive processing, dependencies in immediate versus later processing, and dependencies that are graded versus all-or-none. These subdivisions define eight possible hypotheses. Importantly, these alternatives are not mutually exclusive so if one counts the hybrid models that combine elements of each distinction there are many more possible hypotheses.

In the following section, we present some models of congruency effects combined with our model of limited-capacity, parallel processing. All of the models mediate the congruency effects with selection errors because the interactive processing models predict congruency effects in both single-task and dual-task conditions which were not found. They also all mediate the effect within later processing (memory, decision or response) because in Experiment 3 there were congruency effects in the sequential condition that were as large as found in the simultaneous condition. This is consistent with the role of later processing. This leaves two models to document in detail: a graded model and an all-or-none model.

## A weighting model of congruency effects

In this section we detail a graded model of selection errors. This weighting model can be interpreted as a selection model with graded weights for decision (Kinchla & Collyer, 1974). Alternatively, this model can be used to describe interactive processing such a crosstalk (Navon & Miller, 1987). The equations do not distinguish these hypotheses. To distinguish them, one must depend on the effects of other manipulations such as single versus dual task and the simultaneous versus sequential displays.

We assume all of the features of the previously described limited-capacity, parallel model concerning stimulus representation, target representation, capacity limits, and the decision rule. The change is to add a revision to the decision variable that was *S′*_*1*_ and *S′*_*2*_. Denote a new decision variable for each task as *S″*_*1*_ and *S″*_*2*_. Assume these are a function of the modified stimulus representations *S′*_*1*_ and *S′*_*2*_:

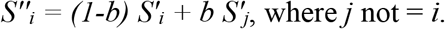

The *selection* parameter *b* is a weight for the inclusion of information from the irrelevant stimulus representation. It is zero for the case of no selection error and is 0.5 for the case of the maximum selection error: equal contributions from both stimulus representations. This form of selection error is motivated to fix the variability of the decision variable to 1. Since both random variables begin with a unit variance, the weighting by *b* and (1-*b*) maintain the variance at 1. Using this addition, the final proportion of “yes” responses is thus given by:

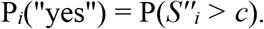

## A mixture model of congruency effects

A mixture model is appropriate when selection fails in an all-or-none way (e.g. Palmer & Moore, 2009). As with the prior model, assume all of the features of the limited-capacity, parallel model concerning stimulus representation, target representation, capacity limits, and the decision rule. The added dependency is that on some trials, the relevant stimulus is replaced by the irrelevant stimulus. As before, the stimuli are represented by random variables *S′*_*1*_ and *S′*_*2*_ for the two tasks (*i* = 1 or 2). Suppose the relevant task (and stimulus) is *i*, the probability of a “yes” response in this task is a mixture of trials in which the decision was based on the relevant stimulus and trials in which it was based on the irrelevant stimulus:

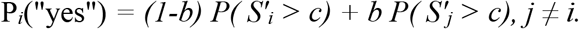

The selection parameter *b* is the probability of basing the decision on the irrelevant stimulus.

## Comparing Predictions of Weighting and Mixture Models

## Congruency effects

We now describe predictions for congruency effects. Begin by considering selection weighting combined with an unlimited-capacity, parallel model (*k* = 0) assuming moderate amounts of selection error (*b* = 0.15). To generate specific predictions, assume equal-variance Gaussian distributions for targets and distractors.

The left panel of Figure 18 shows the congruency effects predicted by the weighting model. A moderate congruency effect is predicted for the dual-task condition and no effect for the single-task condition. For the dual-task and *b*=0.15, the congruency effect is about 8% when the mean performance is 75% correct. As discriminability improves toward perfect, the congruency effect falls back to zero.

**Figure 18.**
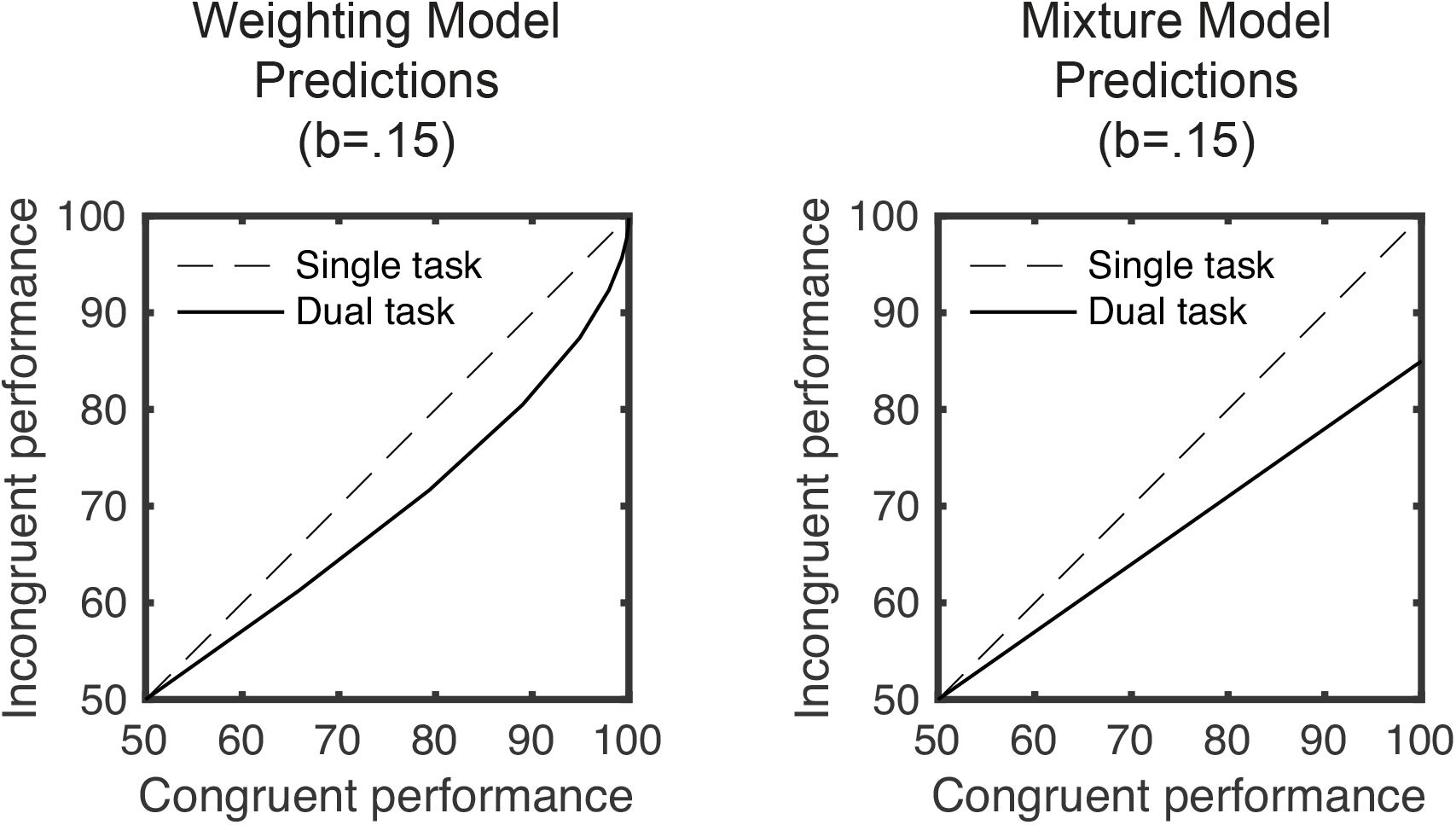
Congruency effects for weighting and mixture models. Two versions of an unlimited-capacity, parallel model are illustrated. For both, the incongruent condition is plotted against the congruent condition. The dashed curve is for the single-task condition and the solid curve is for the dual-task condition. For the weighting model, congruency effects are predicted for the dual-task condition that disappear with high discriminability conditions. For the mixture model, congruency effects are predicted for the dual-task condition that persists even with high discriminability conditions. The results of the Gabor detection experiments are qualitatively consistent with the weighting model.

Next consider the predictions of a mixture model with substitutions of irrelevant stimuli rather than weighting. Again, we assume unlimited capacity (*k=0*), a moderate amount of selection error (*b=.15*), and equal-variance Gaussian distributions.

The right panel of Figure 18 shows the congruency effects predicted by the mixture model. As before there is a moderate congruency effect for the dual-task condition and no effect for the single-task condition. The new feature of the prediction is that the congruency effects grow proportionally with performance. For the dual-task and *b*=0.15, the congruency effect is about 9% when the mean performance is 75% correct. As discriminability improves toward perfect, the congruency effect now grows to a maximum of 15%. The value of this maximum congruency effect is equal to the selection parameter *b*.

## Dual-task deficits

Next consider the dual-task deficits predicted by these two models. The predictions of the weighting model are shown in the left panel of Figure 19. The dual-task deficit is very small and are invisible in this figure. The deficit for a single-task performance of 75% correct is about 0.4%. Thus, adding weighted selection errors causes little change to the dual-task deficit for the two-choice tasks considered here.

**Figure 19.**
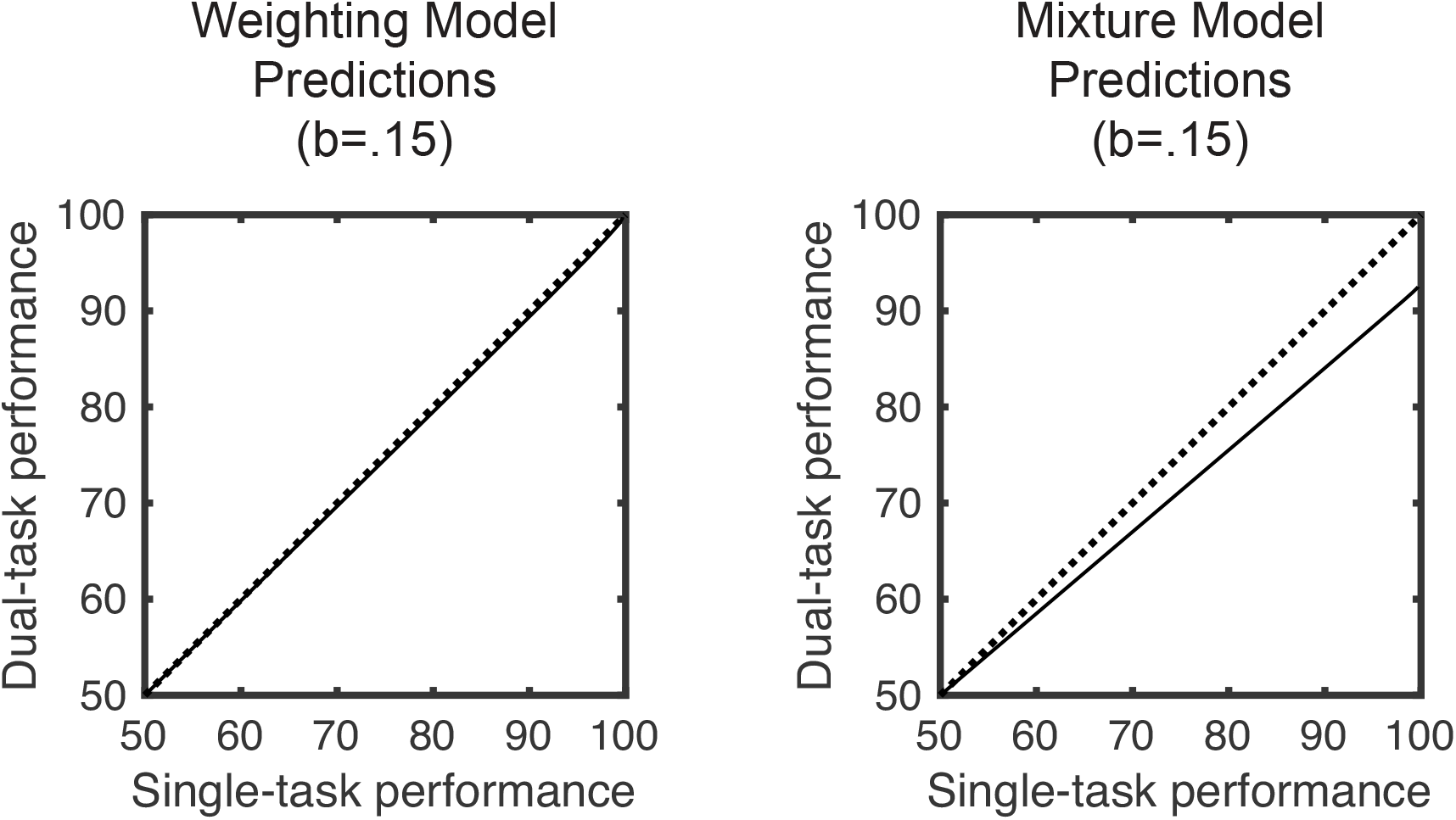
Dual-task deficits for weighting and mixture models. Two versions of an unlimited-capacity parallel model are illustrated. For both, the dual-task condition is plotted against the single-task condition. The dashed curve is for the single-task condition and the solid curve is for the dual-task condition. For the weighting model, there are essentially no dual-task deficits. For the mixture model, there are dual-task deficits that arise from the interactive processing rather than a capacity limit. The results of the Gabor detection experiments are qualitatively consistent with the weighting model.

The right panel of Figure 19 shows the dual-task deficits for the mixture model. Unlike the previous model, adding selection error does change the dual-task deficits. There is now a dual-task deficit even when there is unlimited capacity. For a single-task performance of 75% correct, the dual-task deficit is about 4%. This is because substitution hurts performance in the incongruent condition and cannot help performance in the congruent condition. In short, substitution causes an increase in the dual-task deficit. Thus, a dual-task deficit is a necessary prediction of selection by substitution. The absence of a dual-task deficit is by itself evidence against substitution.

## Response correlations

Next compare the response correlations predicted by the two models. Predictions of the weighting model are shown in the left panel of Figure 20. These predictions are for a dual-task condition with *d*=*2*, *k*=*0* and *b*=*0.15*. We use this intermediate discriminable condition throughout this section. There are positive correlations predicted for the signal-signal and noise-noise trials and negative correlations predicted for the signal-noise trials. The correlation predictions for the case with probability of “yes” equal to .45 are highlighted with a red circle because these are the conditions observed in the experiments. These correlations are not very dependent on discriminability and are similar for *d*=*1*. On the other hand, the correlations are strongly dependent on the selection parameter *b*.

**Figure 20.**
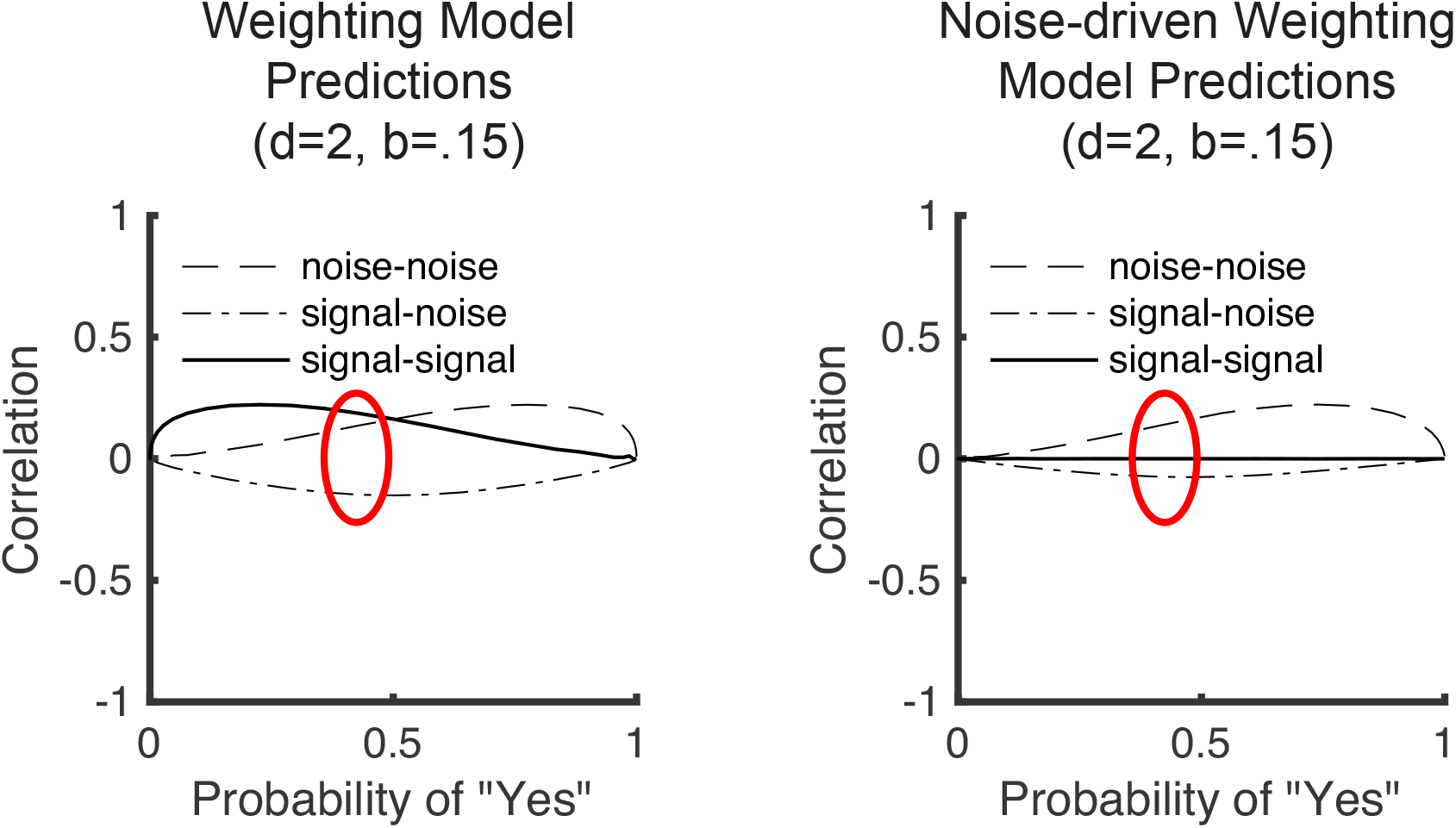
Predicted correlations for two versions of the unlimited-capacity, parallel model. The predicted correlation is plotted against the observed probability of responding “yes”. The three curves are for trials with signal in both stimuli, signal in one and noise in the other, and noise in both stimuli. In both panels, the predictions for p(“yes”) = .45 are circled for emphasis. In the top panel with symmetric selection errors, positive correlations are predicted for signal-signal and noise-noise conditions while negative correlations are predicted for the signal-noise condition. In the bottom panel with noise-specific selection errors, positive correlations are predicted for noise-noise conditions while zero or slightly negative correlations are predicted for the other conditions. The results of the Gabor detection experiments are qualitatively consistent with the noise-selective version of the selection errors.

This predicted pattern does not capture the pattern observed for Gabor detection. In the experiments, the main correlation was positive for only noise-noise trials (see also the color task of White et al., 2018). That pattern can be obtained by changing the model so that there are separate terms for signal and noise conditions. A model with selection weights driven by noise alone is illustrated in in the right panel of Figure 20. Now the positive correlations occur primarily for noise-noise trials. Thus, this noise-specific weighting model results in a pattern of correlation more like that found with Gabor detection.

[Note to coauthors: More can be done here. I want to write the following sentence but am not sure it is true yet: “One process interpretation of this model is that the noise distributions for the two tasks are correlated but the signal distributions are not.”]

The prediction of the mixture model is quite different. We have assumed that the probability of a substitution is independent for the two tasks. This results in a predicted response correlation of zero. Another possibility is the assume that all errors are transpositions. For these two-response tasks, this is equivalent to assuming that substitution errors are perfectly correlated across trials. For the chosen conditions of *d*=*2*, *k*=*0*, *b*=*0.15*, the predicted correlations remain zero for noise-noise and signal-signal trials. But there is a positive correlation of .31 specific to signal-noise trials with a peak at probability “yes” equal to .5 (not shown in the Figures). Neither of these variations yield results similar to what was observed for Gabor detection.

## Summary

The predictions of the weighting and mixture models differ for each of the effects examined here. First, the congruency effect is predicted for highly visible stimuli for the mixture model but not for the weighting model. Second, dual-task deficits always occur for the mixture model but need not occur for the weighting model. Third, different patterns of correlations are predicted by the two models. Only a modified weighting model with noise-specific weights predicts positive correlations specific to the noise-noise condition. Thus, for all three of the effects, one can distinguish the weighting and mixture models. Moreover, the observed results from Gabor detection all favor the weighting model.

## Computational Methods

All numerical predictions were based on simulation. Our goal in these simulations was to achieve at least three digits of accuracy. We confirmed the precision of our estimate making multiple estimates and measuring that the standard deviation of that sample was smaller than 1/1000 of the estimated value. To achieve this goal, 1,000,000 trials were simulated for each predicted proportion. To estimate the response correlations, 10,000,000 pairs of responses were simulated.

To estimate the area under the ROC curve (*A*_*ROC*_), we numerically integrated a stimulated ROC curve. Care was necessary because the area is underestimated if one uses steps that are too large or limits that are not extreme enough. The curve was sampled by varying the criterion in steps of 0.005 and extending the limits of integration to achieve proportions of hits and false alarms that differed from 0 or 1 by less than 0.005. These choices were based on the testing with a range of yet smaller values and picking the largest values that were within 1/1000 of the estimate made with much smaller steps and more extreme limits. Based on these tests, we believe our estimates are good to at least three digits.

## Author Note

We thank xxx and xxx for comments on earlier versions of this article. This research was partially supported by NEI grant EY12925 to Geoffrey M. Boynton.

